# Genetic and structural evidence links Ca^2+^ dysregulation and ATP2B2 to neuropsychiatric illness

**DOI:** 10.1101/2025.08.25.672202

**Authors:** Sherif Gerges, Nikolaj Catois Straarup, Mohamed A. El-Brolosy, F. Kyle Satterstrom, Nolan Kamitaki, Jiayi Yuan, Emi Ling, Carmen Gelze, Raozhou Lin, Melissa Goldman, Curtis Mello, Tarjinder Singh, The Autism Sequencing Consortium, Jonathan S. Weissman, Sabina Berretta, Jen Q. Pan, Hilary Finucane, Charlott Stock, Poul Nissen, Steven A. McCarroll, Mark Daly

## Abstract

Neuropsychiatric disorders are highly heritable, but the molecular mechanisms linking risk variants to disease remain unclear^1^. Linking genetics to biological mechanisms requires integrating evidence across scales, from sequence variation to gene regulation to protein function. Here we integrate genetic, transcriptomic, and structural evidence to identify dysregulation of neuronal Ca^2+^ dynamics as a contributing mechanism. We analyzed single-nucleus neuronal RNA-seq data together with genome-wide association study (GWAS) heritability in a new way to identify gene expression programs enriched for psychiatric risk; genes encoding Ca^2+^ flux pathway genes were implicated by this analysis, a result that was then confirmed by concentrations of rare coding variants in these same genes in persons with psychiatric disorders. A critical gene in this biology, *ATP2B2*, encodes a Ca^2+^-extruding ATPase pump^2^ linked to neuropsychiatric disorders primarily through missense variants. To recognize specific molecular functions affected by these missense variants, we developed a 3D-neighborhood-based method that maps missense variants onto AlphaFold3-predicted protein structures and tests clustering of protein-altering variants in three-dimensional space. This approach revealed clustering of these missense changes in the Ca^2+^ pore and ATP:Mg^2+^ coordination site of ATP2B2. To validate the structural predictions arising from the genetic analysis, we determined the structure of human Ca^2+^-bound ATP2B2 at 2.64 Å resolution by cryogenic electron microscopy. The structure replicated the 3D mutational hotspots identified by the genetic analysis, supporting the observation that neuropsychiatric variants cluster near specific catalytic sites of ATP2B2. In vitro experiments revealed that missense variants prioritized by the structural clustering analysis impaired ATP2B2-mediated Ca^2+^ extrusion in cellular and biochemical assays. Together, these findings suggest that precise regulation of Ca^2+^ dynamics is a contributing mechanism in neuropsychiatric disorders and establish a structure-based framework for predicting the mechanistic impact of missense variants.

## Introduction

Neuropsychiatric disorders impose a substantial global burden ^3^, yet decades of drug development have yielded few novel mechanisms of action^4–7^. Human genetic studies have identified hundreds of loci ^8–11^, but translating these into neurobiological insights remains challenging: most common variants have small effects and lie in noncoding regions, obscuring causal genes and biology^1,12^.

Protein-coding variants can directly implicate specific genes, yet translating their patterns into mechanistic understanding remains a central challenge. Risk-increasing coding variants are typically rare or ultra-rare^13–15^, and the vast majority of coding variants are missense, which present two challenges: first, their effects lie along a continuum from benign to damaging, and it is generally unclear which missense variants are truly pathogenic; second, their mechanistic consequences are highly diverse and challenging to predict computationally^16^. Furthermore, pathogenicity predictors (often based on e.g. conservation or variation) exhibit variable performance across genes and are vague about mechanism, whereas experimental validation remains challenging to scale to many variants^17–20^. These challenges are amplified in the setting of the human brain, which is unrivaled in its organized complexity^21,22^.

To identify molecular programs through which common variant risk converges in neuropsychiatric disorders, we focused on neurons, the cell class most often implicated across human genetic and functional studies, which consistently recognize synapses as a locus for large minorities of these genetic effects ^8,14,23–25^. We analyzed transcriptomic covariation across neuronal cell types spanning multiple regions of the human brain in a way that integrated these expression relationships with per-SNP heritability from genome-wide association studies (GWAS) (**Figure 1a**). This analysis highlighted Ca^2+^ flux pathway genes^26–28^ — channels, pumps, and exchangers that shape the magnitude, duration, and shape of Ca^2+^ fluxes — as one of the strongest functional domains of risk.

**Figure 1.**
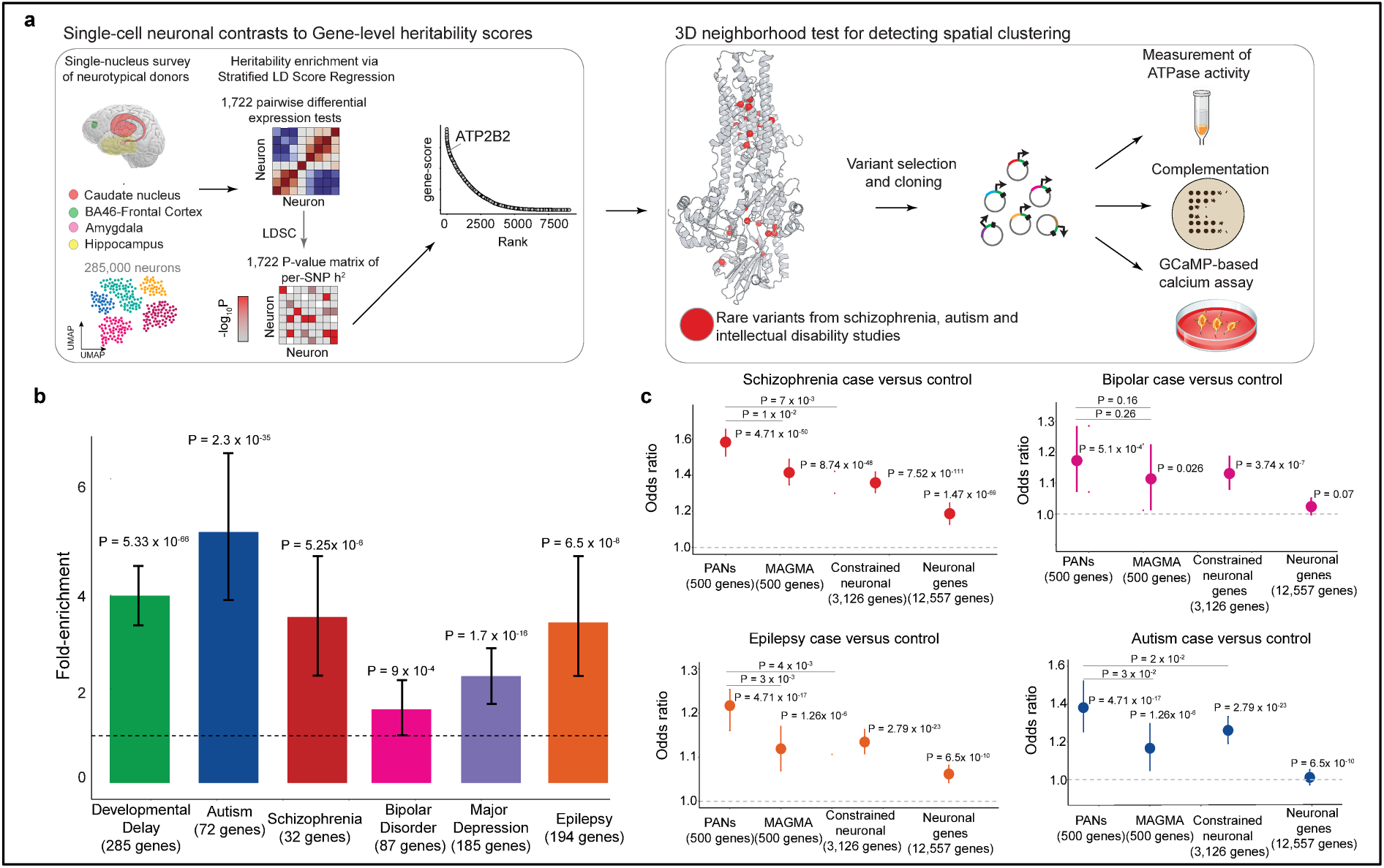
Overview. A. We performed a pairwise differential expression analysis, integrating snRNA-seq with GWAS summary statistics to identify gene sets whose coordinated expression patterns are enriched for common variant heritability. The framework performs bidirectional pairwise differential expression across neuronal cell types, tests each resulting gene set for heritability enrichment using stratified LD score regression, and aggregates significant contrasts into gene level scores and a heritability weighted co-occurrence network. B. Results demonstrating fold-enrichment of prioritized gene scores in exome sequencing data for neurodevelopmental and psychiatric disorders. Error bars represent the 95% CI from a bootstrap of the prioritization scores for significant versus non-significant genes in each phenotype (Supplementary Methods). P-values are from a wilcoxon rank sum test between significant and non-significant genes. For autism, developmental disorder, and schizophrenia we use genes denoted by their significance threshold in the original publications (FDR ≤0.001, Bonferroni significant, and FDR ≤0.05, respectively). For bipolar disorder, major depression, and epilepsy, we use genes with either a protein-truncating or missense p-value < 0.01. C. Case-control enrichment of ultra-rare coding variants from the SCHEMA consortium (n=22,444 cases and n = 39,837 controls) compared to the top 500 constrained genes prioritized by MAGMA, as well as global lists of genes derived from combining constraint and expression. The reported p value was obtained from a gene set burden test in exome data. We counted protein-truncating variants and damaging missense variants with MPC greater than 3 in cases and controls across the top 500 constrained prioritized genes, aggregated variant counts across genes, accounted for sample sizes, and compared cases and controls using Fisher’s exact test on a 2×2 contingency table. The p-values are derived from permutation testing (n = 10,000) of prioritized genes against random constrained genes selected from the full MAGMA list for genes with a Z-score > 0.

Within this set, ATP2B2 stood out for a genetic signal (ultra-rare mutations in psychiatric patients) driven predominantly by missense variants, in contrast to many risk genes enriched for protein-truncating variants. The prominence of missense variants in ATP2B2 suggested disruption of specific structural features. Prior work has shown that disease-associated missense variants can cluster in protein structure^29^, especially in Mendelian disease, cancer ^30–32^, and neurodevelopmental disorders^33,34^. Variant effect predictors like AlphaMissense^17^, PopEVE^35^, deMAG ^36^ do not identify clusters. A systematic case-control framework for identifying disease-enriched 3D neighborhoods has been lacking.

We developed the 3D neighborhood test (3DNT), which integrates either experimentally determined or AlphaFold3-predicted^37^ protein structures with any dataset of missense variants ascertained in cases and controls. Applying this approach to rare missense variants from schizophrenia exome sequencing (SCHEMA)^14^ and *de novo* variants from autism spectrum disorder (ASD) and neurodevelopmental disorders (NDDs)^38^. The analysis identified discrete 3D neighborhoods of missense variants within ATP2B2, from which we selected representative variants for functional assays of calcium handling and ATPase activity.

## Results

### Pairwise analysis of neuronal gene expression

We first developed a computational approach for gene prioritization that exploited information about the diverse gene-expression modules and programs that are utilized to variable extents by diverse types of neurons. The basic idea of this approach was to use the information in gene co-expression: recognizing sets of genes that are consistently co-regulated with genes that are enriched for genetic effects on risk of psychiatric disorders. We used single-nucleus RNA-sequencing (snRNA-seq) data from 42 neuronal cell types sampled from multiple human brain regions (dorsolateral prefrontal cortex^23^, caudate^39^, amygdala and temporal lobe), integrating these data with GWAS summary statistics (association statistics for SNPs in genes across the genome) to identify genes that tended to be co-regulated with sets of genes enriched for common genetic influences on risk of psychiatric disorders (**Figure 1**). To do this, for each cell-type pair, we performed a bidirectional pairwise differential expression test, identifying genes more highly expressed in one neuronal cell type relative to the other, and vice versa, thereby generating a large collection of gene sets defined by these pairwise comparisons. We then tested each such set (1,766 sets) for enrichment of common variants that contribute to the heritability of neuropsychiatric disorders^40–42^. This approach yielded a per-gene prioritization score, which was computed as the sum of heritability enrichment z-scores (*t*/standard error) over all FWER-significant gene sets containing the gene.

When prioritized using the schizophrenia GWAS data, scores were assigned to 9,100 of 17,468 genes (Supplementary Table 7), then used to rank genes in the analyses below; the remaining genes were not assigned a score, as they did not appear in any pairwise gene set showing significant heritability enrichment by LDSC (see Methods and Extended Discussion Note).

### Prioritized genes capture biologically meaningful rare variant signal

Since these genes had been prioritized by their relationships to common genetic variation associated with schizophrenia, we critically evaluated these gene rankings using a different kind of human genetic data – ultra-rare (minor allele frequency < 0.01%) protein-coding variants ascertained in persons with neuropsychiatric disorders; this validation approach leverages the convergence but independence of common and ultra-rare variant risk on shared genes^43,44^. Of constrained genes (LOEUF decile < 2), the 500 genes most strongly prioritized by this analysis were significantly enriched for rare coding variation across multiple neuropsychiatric disorders and showed stronger signal than genes prioritized by existing methods using GWAS data or using gene-expression data alone ^45^ (**Figure 1b, 1c, S2b**, Methods).

We then tested the 500 most prioritized genes for enrichment of psychiatric drug targets (largely identified independently of GWAS studies)^6,46^. Curated drug target genes from Open Targets showed enrichment of rare coding variation in exome data for multiple disorders (**Figure S2c**), including several ion channel and synaptic genes relevant to neuronal signaling, suggesting that the integration of transcriptome and GWAS results had prioritized genes with biological relevance. The results in schizophrenia were strongest statistically (likely owing to availability of extremely well-powered GWAS and exome sequencing studies) and became the primary focus of downstream analyses.

### Rare variant burden in calcium flux genes is enriched across neuropsychiatric disorders

To identify functionally related sets of genes implicated by these GWAS/transcriptome gene rankings, we tested the top 500 prioritized genes for schizophrenia for enrichment in gene ontology categories^47,48^. The most significant signals consistently converged on calcium signaling, including regulation of postsynaptic cytosolic calcium and intracellular Ca^2+^ levels (**Figure 2a**). Similar patterns were observed across additional brain-related traits (**Figure S3a**, see **Supplementary Table 3** for the full list of gene ontology enrichments).

**Figure 2.**
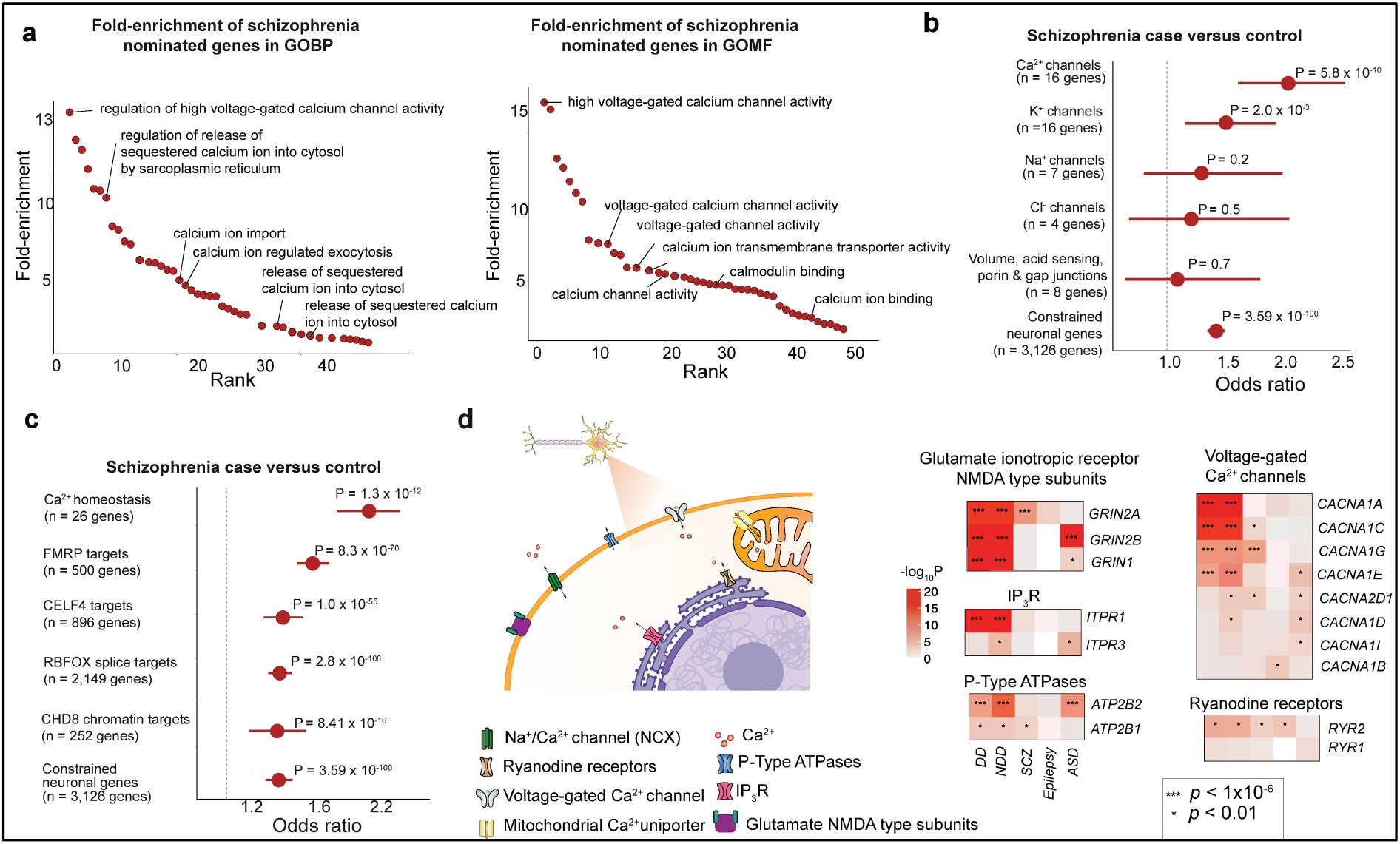
Ca^2+^ flux genes are enriched for ultra-rare protein-coding variants in schizophrenia. A. Fold enrichment of the top 500 schizophrenia-nominated genes in Gene Ontology Biological Process (GOBP) and Gene Ontology Molecular Function (GOMF) terms. The x-axis represents the rank of the GOBP terms based on their fold enrichment; the y-axis shows the fold enrichment values. Each point corresponds to a GOBP/GOMF term. The highest fold enrichment is observed in terms related to calcium ion channel processes. B. Left: Case-control enrichment of ultra-rare protein-coding variants associated with schizophrenia in major voltage-gated ion channels genes. The two-sided P values from Fisher’s exact test displayed were obtained by comparing the burden of variants of the labeled consequence in cases and controls. The dot represents the OR and the bar represents the 95% Cl of the point estimates. Numerical results are in Table S1. C. Case-control enrichment of ultra-rare protein-coding variants from SCHEMA in constrained gene members within the Ca^2+^ flux pathway genes compared with functional neuronal and regulatory gene sets (the number of genes are shown in parentheses below each set). The two-sided P values from Fisher’s exact test displayed were obtained by comparing the burden of variants of the labeled consequence in cases and controls. The dot represents the odds ratio and the bar represents the 95% CI of the point estimates. D. Right: Schematic of Ca^2+^ flux pathway genes. Genes with exome-wide significance in any phenotype are highlighted in red, nominally significant genes are highlighted in blue. Left: Heatmaps of p-value associations in exome sequencing studies of neuropsychiatric disorders of select members of this set of genes. [Figure created with BioRender.com]

Genetic studies of neuropsychiatric disorders have previously implicated voltage-gated ion channel genes^14,49,50–53^. To determine whether this pattern has an underlying specificity to a specific ion (rather than a general property of ion channels), we stratified voltage-gated channels by ion selectivity (Ca^2+^, K^+^, Na^+^, CI^−^; see Methods: Ion Channel Gene Set Selection)^54^. We performed a burden test of PTVs and damaging missense variants (MPC > 3) from schizophrenia cases and controls across these gene sets, and observed the strongest enrichment in Ca^2+^ voltage-gated channels (Fisher’s exact test OR = 1.55, P = 5.52 × 10^−10^)—a rate substantially exceeding that of other channel classes (**Figure S4a**). A similar pattern was observed in ASD, for which Ca^2+^ voltage-gated channels also showed the strongest enrichment. We did not observe such a pattern for genes implicated in epilepsy, consistent with its genetic architecture being dominated by coding variation in sodium channels^55^. (Sodium channels tend to be the fast, high-density drivers of the action potential upstroke, contributing substantially to neuronal excitability and seizures, while calcium often acts as a second messenger that couples electrical activity to gene transcription, synaptic plasticity, and activation of specific enzymes downstream of synaptic activity.) These results indicate that calcium, more than other ionic substrates, is the driver of the genetic signal that arises from voltage-gated ion channel genes in multiple neuropsychiatric disorders.

We then asked whether these results reflect a broader principle of calcium transport rather than being specific to channels. To test this, we systematically identified all protein-coding genes involved in ion transport using HGNC-curated gene sets, spanning both presynaptic and postsynaptic compartments, as well as intracellular organelles and non-selective/other (including volume-regulated, acid-sensing, porin, and gap junction channels). Strikingly, the Ca^2+^ “superset” of channels exhibited the strongest enrichment relative to all other ion channel classes (OR = 1.99; P = 5.85×10^−10^, **Figure 2b**). Notably, the prioritization also highlighted calcium-signaling genes with emerging genetic implication from rare protein-coding genetic variants, including *RYR2*, an endoplasmic reticulum (ER) calcium release channel (SCHEMA P = 3.4×10^−3^; epilepsy P = 2.8×10^−4^; NDD P = 2.86×10^−6^), and *ITPR1*, an inositol trisphosphate receptor mediating release of calcium from the ER (NDD P < 2.2×10^−16^). The prioritization further implicated genes involved in calcium transport that fall outside the HGNC ion channel classification, including *SLC8A1*, a sodium-calcium exchanger. This prompted us to curate a more comprehensive Ca^2+^ channel gene set encompassing all genes encoding proteins that enable calcium to move across membranes.

These genes, which we collectively referred to as Ca^2+^ flux pathway genes ^27,56–61^, (CFPG hereafter) encompass channels, pumps, and exchangers that orchestrate spatial and temporal control of calcium in a cell and its various compartments such as dendrites and synapses (full list in **Supplementary Table 4**). We compared constrained members of this CFPG gene set to other nervous system gene sets with strong associations to schizophrenia, namely translational targets of FMRP^62^, chromatin targets of CHD8^63^, and splice targets of RBFOX^64^, and found that CFPG showed significantly greater enrichment for rare variants in persons with schizophrenia (OR = 1.96 and P = 1.3×10^−12^, **Figure 2c**). We examined *de novo* variants, analyzing data from the 2026 Autism Sequencing Consortium cohort (manuscript in preparation, 38,680 ASD probands and 9,567 non-ASD siblings) and NDD. Across both datasets, constrained CFPGs showed disproportionately high *de novo* missense rate ratios compared to other nervous system gene sets, while PTV rate ratios were comparatively modest—both relative to their own missense signal and relative to PTVs in other gene sets (Figure S4b). This missense-biased signature may reflect the sensitivity of Ca^2+^ dynamics in the cell, where even modest perturbations can disrupt sensitive calcium dynamics and the downstream pathways they instruct ^57,65^.

Among the most highly prioritized genes genome-wide were *ATP2B1* (ranked #8) and *ATP2B2* (ranked #12), members of the plasma membrane Ca^2+^-ATPase (PMCA) family (known as PMCA1 and PMCA2), which extrude calcium and thereby regulate calcium levels^66^ at the sites where they are localized, including dendritic spines and presynaptic terminals^67^. The prominence of *ATP2B1* and *ATP2B2* suggests a role for calcium clearance^67^ – in addition to the calcium influx mediated by voltage-gated calcium channels – suggesting the importance of precise control of calcium dynamics in these subcellular structures.

Furthermore, both *ATP2B1* and *ATP2B2* show compelling genetic evidence of involvement in neuropsychiatric disorders. Mutations in *ATP2B1* are an established cause of NDDs^68^; *ATP2B1* also carries a nominally significant burden of PTVs and damaging missense variants (determined by Missense badness, PolyPhen-2, and Constraint, MPC^69^) in schizophrenia (SCHEMA PTV and MPC > 3, P = 8.7 × 10^−3^). *ATP2B2* harbors a missense-specific schizophrenia signal (MPC 2–3 OR = 2.0, P = 7.0 × 10^−4^) which, while not exome-wide significant, ranked 5th in SCHEMA in missense burden and was more recently implicated in a large Utah pedigree^70^. *ATP2B2* is exome-wide significant in both ASD and NDD (q < 5 × 10^−6^ for both disorders) ^38,71,72^ and, similar to the missense-specific schizophrenia signal, 19 *de novo* missense mutations and only a single protein-truncating mutation have been observed across ASD and DD/ID studies. This unusual pattern of missense-driven risk suggested a potential opportunity to identify the specific aspects of ATP2B2 molecular function that are important for protection from schizophrenia, as well as an avenue for refining and better understanding the reasons that constellations of missense variants appear more frequently in cases than controls – a key challenge in schizophrenia and indeed in all of human genetics.

### 3D neighborhood analysis highlights key functional elements

The excess of missense variation affecting ATP2B2 in schizophrenia and autism could in principle involve the selective perturbation of specific functional domains. To test this, we developed a molecular-structure-based analytical framework that recognized the clustering of missense variants in the encoded protein’s three-dimensional structure; we call this approach the “3D neighborhood test”. We aggregated missense variants from SCHEMA together with *de novo* missense variants from ASD and NDD cohorts, assigning “case” status to variants observed only in cases/probands and “control” status to variants observed only in controls. We mapped all 331 variants onto the AlphaFold3-predicted structure of ATP2B2 and restricted analysis to structurally well-modeled regions (see Methods). For each residue harboring one or more variants, we defined a three-dimensional neighborhood of radius 15 Å and tested for enrichment of case versus control variants relative to the rest of the protein using Fisher’s exact test (Figure 3a; Supplementary Table 9 for list of p-values).

**Figure 3:**
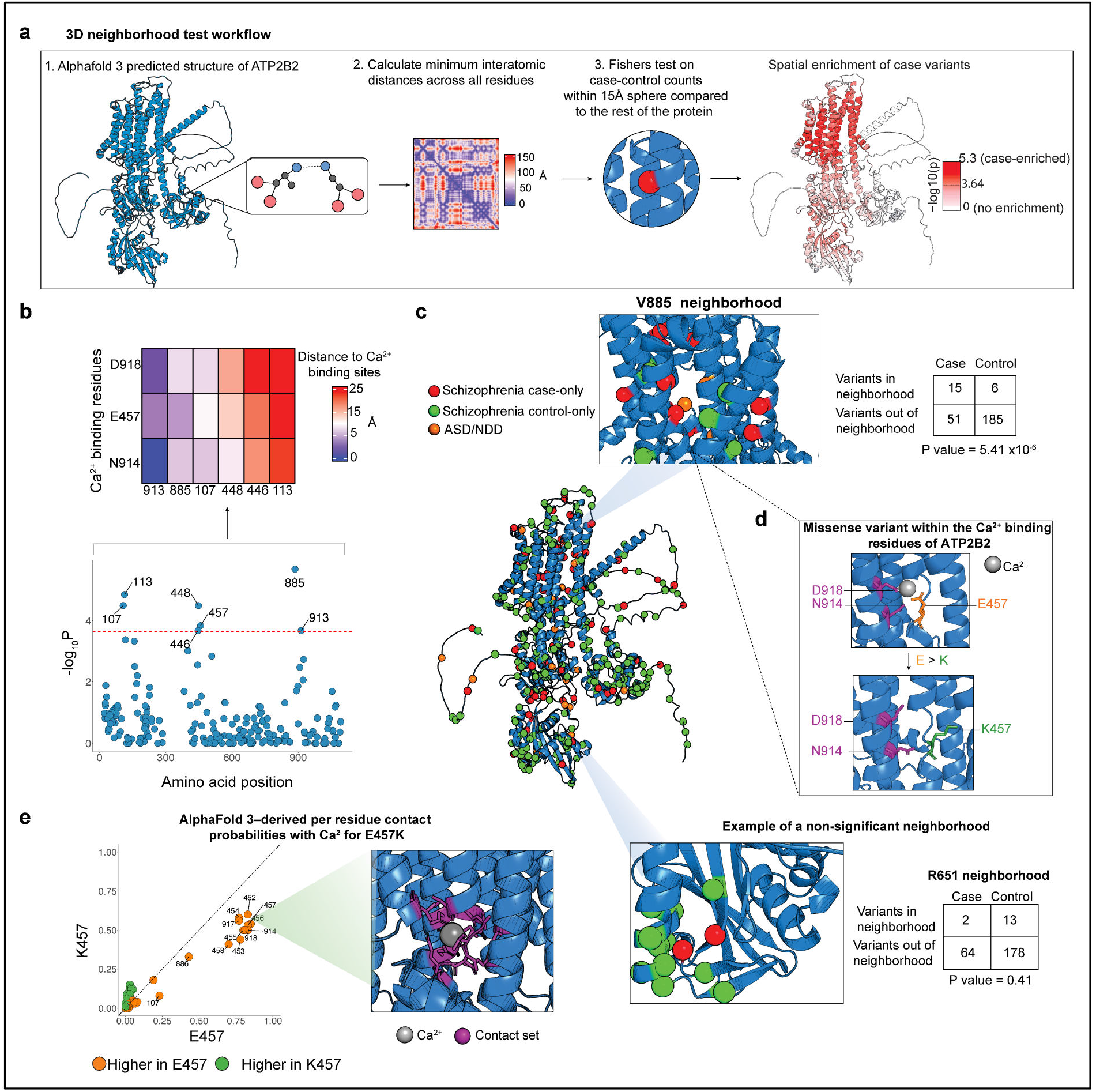
3D-neighborhood analysis of neuropsychiatric-associated variants reveals impact on ATP2B2 Ca^2+^ binding and activity. A. Outline of the 3D neighborhood method. Using the AlphaFold3 structure of ATP2B2, we computed the shortest interatomic distances between all pairs of residues after excluding regions with low pLDDT and high PAE. For each residue, we defined a sphere of radius 15 Å and used Fisher’s exact test to assess enrichment of case or *de novo* missense variants associated with schizophrenia, autism, and NDD within the sphere compared to the remainder of the protein. On the far left, Each residue is colored by −log10 of the smallest Fisher’s exact p-value across all 15 A neighborhoods containing it. Dashed line on color bar marks the Bonferroni significance threshold (p = 0.05/219 tested residues). B. All significant residues are in or around the Ca^2+^-pore of ATP2B2. Bottom: P-values from the per-residue neighborhood burden enrichment analysis. The red line represents the Bonferroni significance threshold, with labeled residues indicating those reaching statistical significance. Top: Heatmap of the distances (in A) of each of the Bonferroni significant residues to the three principal Ca^2+^-binding sites in *ATP2B2*. C. Case-only schizophrenia, ASD, and NDD-associated variants are mapped onto the AlphaFold-predicted structure of *ATP2B2*. Highlighted are two regions from the result of the per-residue burden analysis (top: residue V885, bottom: residue R651). 2×2 tables adjacent to the plot indicate the case-control counts from the neighborhood analysis corresponding to that highlighted residue. The magenta residue indicates a residue is mutated in both schizophrenia and ASD. D. Ca^2+^-binding site in *ATP2B2* (PMCA2) illustrating the Ca^2+^ binding residues in transmembrane domains IV and VI. The E457K variant disrupts one of the three Ca^2+^ binding residues in the channel. In purple are the two other binding residues, D918 and N914. E. Per-residue calcium-binding probabilities across ATP2B2 comparing E457 and K457. Left: Visualization of the residues most impacted.

We identified multiple 3D neighborhoods that were significantly enriched for “case variants” (variants ascertained in persons with these disorders) after Bonferroni correction. Both *de novo* mutations and variants analyzed by case/control comparisons clustered in the same structural neighborhoods. The strongest signal arose from a neighborhood centered on V885I (OR = 8.97, 95% CI 3.10–29.70, P = 5.4 × 10^−6^; **Figure 3b**), a missense variant observed in schizophrenia cases. Some 22.7% of all case and *de novo* variants localized to this region, in comparison to only 3.1% of control variants, corresponding to a ~7-fold enrichment. To determine whether these neighborhoods are important to ATP2B2 function, we aligned the AlphaFold3 ATP2B2 model to the experimentally resolved cryogenic electron microscopy (cryo-EM) structure of human ATP2B1^73^ (Figure S11a). The V885I-centered neighborhood, along with several additional significant neighborhoods, encompassed the canonical Ca^2+^-binding site (Figure 3d), suggesting that the disruption of calcium binding and transport is the dominant source of signal. To rule out chance, we permuted case and control labels across variants while preserving variant positions and re-ran the scan; the V885I-centered cluster was more extreme than the minimum p-value of every one of 1,000 permuted scans (P = 6 × 10^−3^; 1,000 permutations, Fig. S5d). A second enriched neighborhood centered on E457K, itself a Ca^2+^-coordinating residue and located 11.0 Å from V885I (OR = 6.47, P = 2.2 × 10^−5^; **Figure 3d**). The pathogenicity of this variant is supported by multiple lines of evidence. E457K has been observed independently as a *de novo* mutation in two neurodevelopmental disorder probands. In a mouse model, E457K reduces the rate of intracellular Ca^2+^ increase during neuronal depolarization^74^. In the Regeneron Genetics Center Million Exome dataset, E457 is the second most missense-constrained residue in the entire gene (**Figure S5b**)^13^. E457 is conserved in all Ca^2+^- and Na^+^,K^+^-ATPases (ATP1, ATP2 families) and plays a key role in gating ^75^.

E457K results in charge reversal of the residue, the equivalent mutation in Sarco-Endoplasmic Reticulum Ca2+-ATPase (SERCA, ATP2A1 gene – E309K mutation) blocks calcium binding^76^, and we assumed that PMCA function would be similarly disrupted. We used AlphaFold3 to estimate per-residue Ca^2+^ contact probabilities in wild-type ATP2B2 and ATP2B2-E457K with a single Ca^2+^ (see Extended Discussion: *Simulating Mutation Effects on Per-Residue Contact Probabilities with AlphaFold3* and Figure S6), revealing a marked reduction in Ca^2+^ contact probabilities with residues along the channel pore in the mutant protein (**Figure 3e**). Consistent with these results, Missense3D^77^ predicted that E457K expands the Ca^2+^-binding cavity volume (**Figure 3d**).

While E457K disrupts Ca^2+^ coordination directly, other case variants introduce similar physicochemical disruptions. For example, A400E (observed in one schizophrenia case) and R1014C (observed in one ASD case) alter charge at residues lining the transmembrane pathway, introducing electrostatic changes within a membrane-embedded environment optimized for charge neutralization and hydrophobic packing^78,79^. Other substitutions shift polarity within the transport pathway (S922L, P460S, I1070T, and I966M), replacing polar residues with hydrophobic ones or vice versa. A further group (I446V, V448M, A907G, V913M, V1036M, and F1040L) modifies side-chain volume. Although chemically diverse, these substitutions accumulate around a physically constrained and electrostatically tuned transport pathway, consistent with destabilization of Ca^2+^ binding and changes in local features that may impair ion translocation.

### Conditional analysis identifies a second mechanism of catalytic disruption in ATP2B2

To test for independent neighborhood signals outside the Ca^2+^ pore, we performed a conditional analysis in which we excluded all variants from the primary 3D neighborhood centered on V885I and repeated the test. This revealed a single Bonferroni-significant neighborhood centered on residue R508 (P = 2.01 × 10^−10^; **Figure 4a**), distant from the Ca^2+^-binding site but proximal to the ATP:Mg^2+^-binding pocket (**Figure 4b**). This second signal mapped to the enzyme’s catalytic core, near the ATP-binding interface that powers the transport cycle. After conditioning on the first signal by excluding all variants within the first neighborhood, this region captured 26.0% (N = 12/46) of residues with case variants compared to only 6% (9/148) of control residues, corresponding to an approximately sixfold enrichment (OR = 6.06, P = 2.75 × 1^0−5^).

**Figure 4:**
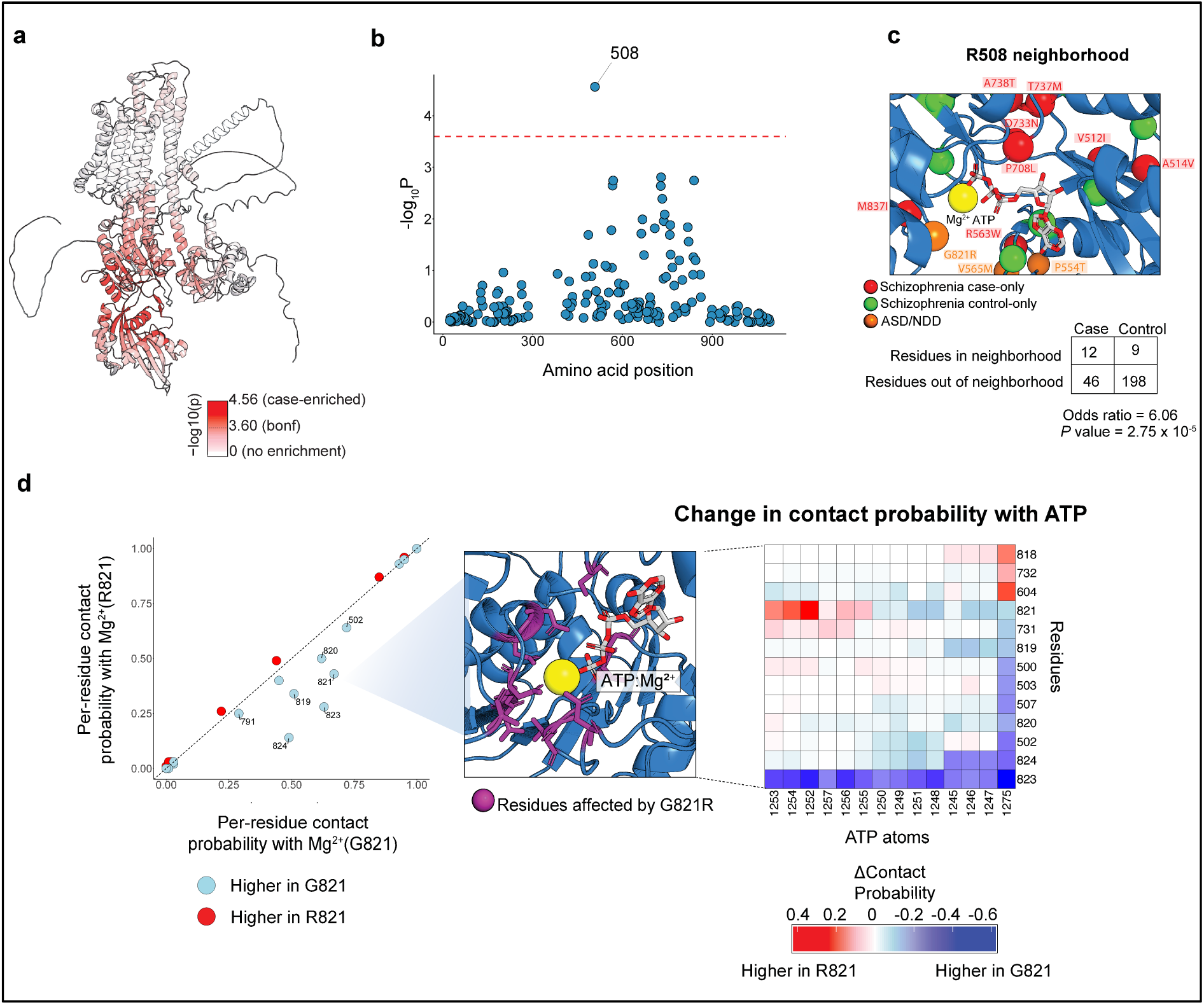
Conditional analysis implicates ATP: Mg^2+^ coordination in ATP2B2 dysfunction. A. Spatial enrichment of case variants across ATP2B2 (conditional analysis). Each residue is colored by −log10 of the smallest Fisher’s exact p-value across all 15 A neighborhoods containing it, after excluding case variants in the primary calcium-binding hotspot. A secondary enrichment emerges in the ATP-binding domain. Dashed line on color bar marks the Bonferroni significance threshold. B. Conditioning on variants clustered at the Ca^2+^ binding site revealed an additional, independent signal at the ATP: Mg^2+^-binding site, suggesting that missense variation may also impair ATPase activity through disrupted ATP hydrolysis. The panel shows P-values from this conditional analysis, with the red line marking the Bonferroni significance P-values (similar to Figure 3B) of the conditional analysis. The red line represents the Bonferroni significance threshold. C. Case-only schizophrenia, ASD, and NDD-associated variants are mapped onto the AlphaFold-predicted structure of ATP2B2 around the ATP:Mg^2+^ binding site. 2×2 tables adjacent to the plot indicate the case-control counts from the neighborhood analysis corresponding to that highlighted residue. D. The substitution G821R modifies the per-residue contact probabilities of residues near the Mg2+ binding site (left) and the ATP binding site (heatmap: right), as computed using AlphaFold3. Middle: Visualization of the residues most impacted by G821R.

Because ATP binding and hydrolysis at this interface powers Ca^2+^ extrusion, spatial enrichment of case variants in this region implicates a second pathogenic mechanism distinct from Ca^2+^ binding itself: disruption of the Post-Albers cycle^80^, specifically by perturbing the binding of the ATP:Mg^2+^ cofactor. Notably, the variant G821R is in closest proximity (within 4.24 Å of Mg^2+^) to the ATP:Mg^2+^ cofactor complex (Figure 4b). Consistent with our analysis of E457K, ligand-protein contact probability modeling of the G821R variant revealed reduced contact probabilities with the ATP:Mg^2+^ complex (Figure 4c), suggesting that G821R may destabilize nucleotide and cofactor interactions critical for E1-E2 transitions and pump function in ATP2B2. Other case variants introduce additional perturbations to nucleotide-dependent function, including D733N (seen in one schizophrenia case), and R588H and R563W, which attenuate charge. Other variants shift polarity, such as T737M and A738T, which alter hydrogen-bonding potential, and V565M, M837I, V512I, A514V, and P708L, which modify side-chain volume.

### Human PMCA2 cryo-EM structure

Motivated by the structural clustering observed in AlphaFold3 models, we sought to directly evaluate these signals in an experimentally determined structure of human ATP2B2. No high-resolution structure of human ATP2B2 has previously been determined, despite its implication with several diseases^81–83^, and while AlphaFold3 provides valuable predictions, it has limitations in capturing conformational and ligand-bound states^84^.

We reconstituted human ATP2B2 (isoform PMCA2z/a) into Salipro nanoparticles and determined its structure by single-particle cryo-electron microscopy in the calcium-bound E1 state at 2.64 Å resolution (Figure 5a, 5b) (Figure S13; Methods). We then re-ran the 3D neighborhood test on this experimentally determined structure. We found eight Bonferroni-significant centers that defined neighborhoods covering 28.3% of residues (272/960), yet captured 39.1% of case variants (25/64) compared to 12.6% of control variants (24/191), a 4.46-fold enrichment. To test for independent signals, we performed a conditional analysis excluding all variants within 15 Å of these neighborhoods; no additional neighborhoods reached Bonferroni significance, although the top residue remained the same as in the conditional AlphaFold3 analysis described earlier. Altogether, these results mirrored those obtained from AlphaFold3 models (Pearson r = 0.97, two-sided P = 1.199 × 10^−152^, Figure 5c).

**Figure 5:**
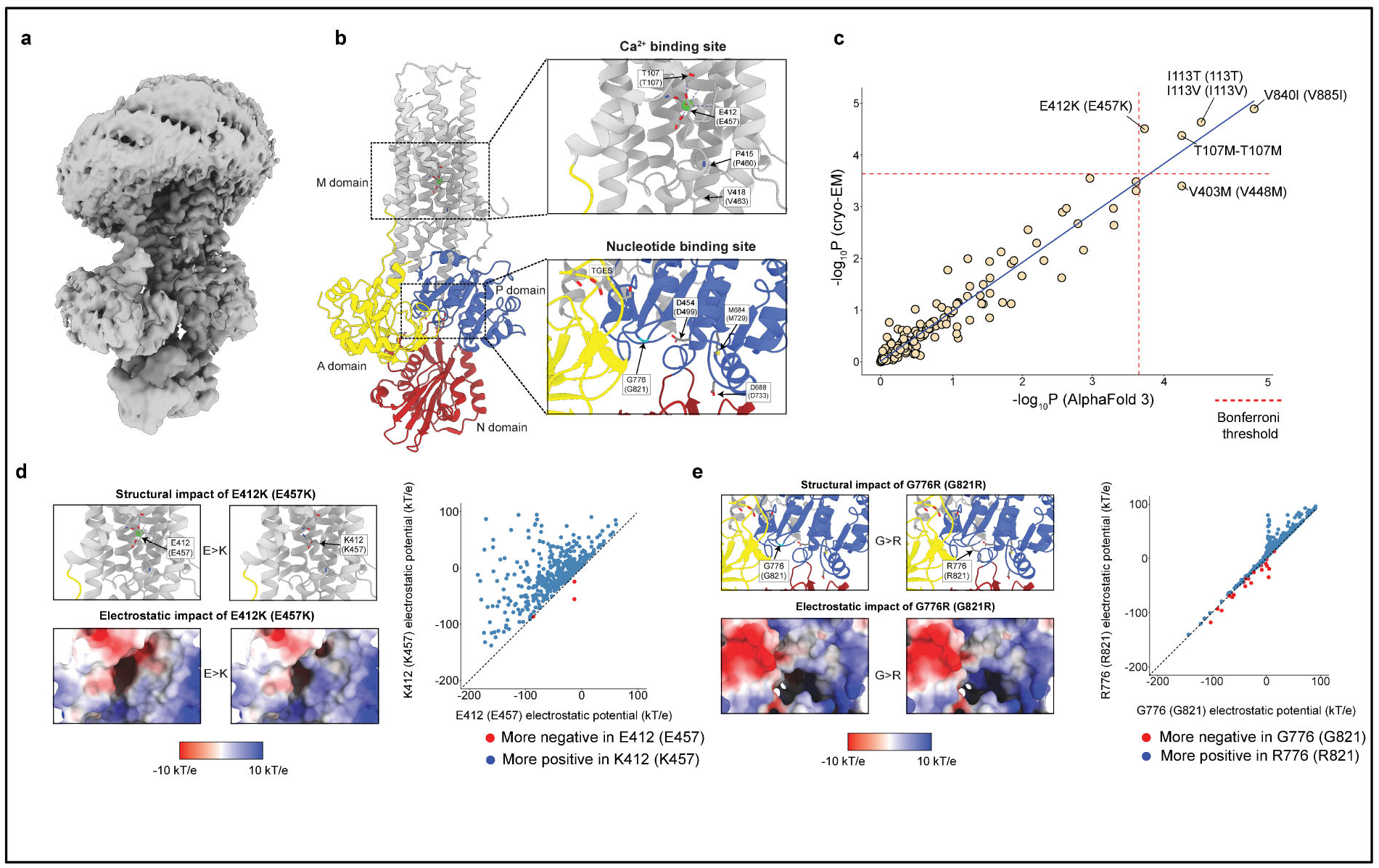
Cryo-EM structure of human ATP2B2 (isoform PMCA2z/a) in an E1 Ca^2+^-bound state. A. Cryo-EM map of human PMCA2z/a. B. Structure with nucleotide binding (N), phosphorylation (P), and actuator (A) domains in red, blue, and yellow, as well as the 10-helix transmembrane (M) domain in silver. Boxes mark the zoomed-in ion and nucleotide binding sites. Mutated residues (T107M, E412K, P415S, V418A, D454N, M684V, D688N, and G776R) are shown in stick representation. TGES dephosphorylation loop is far from the conserved D454 indicating an E1 state. C. Correlation of 3D neighborhood significance values derived from cryo-EM structure and AlphaFold3 prediction. The number before the dash denotes the residue position in canonical ATP2B2, and the number after denotes the corresponding position in the PMCA2z/a isoform. In all figures, splice variant z positions are indicated without brackets, while UniProt positions corresponding to the splice variant w are shown in brackets. D. Top, structural representation of E412K (E457K). Bottom, representation of electrostatic surface potentials computed by APBS for wild-type E412 and the K412 mutant are shown on the protein surface, colored by electrostatic Coulomb potential ranging from-10 kcal (mol e) (red) to +10 kcal (mol e) (blue). Right, the E412K substitution produces a strong local shift from negative to positive electrostatic potential. Shown is the pointwise comparison of electrostatic potentials within a 7 A neighborhood of residue 412. E. Same as D, but for G776R (G821R).

The ligand-protein contact probability modeling indicated that the E457K and G821R substitutions reduced calcium permeability and ATP binding, respectively, which we attributed to local electrostatic perturbations arising from a charge reversal at residue 457 and the introduction of a positive charge at 821. E457 corresponds to E412 in the PMCA2z/a cryo-EM structure, and G821 corresponds to G776. To interrogate the structural consequences of these substitutions, we introduced the E412K and G776R mutations in silico using FoldX|^59^ and computed electrostatic surface potentials using the Adaptive Poisson-Boltzmann Solver (APBS)^85,86,87^ (Methods: Electrostatic potential calculations). We quantified substitution-induced changes in electrostatic potential within a 7 Å sphere centered on each residue, providing spatially resolved support for the predicted functional consequences of each variant. For E412K, the maps confirmed a reversal of surface potential at the Ca^2+^ binding site (**Figure 5e**); for G776R, electrostatic redistribution in the ATP:Mg^2+^ binding region corroborated impaired nucleotide coordination (**Figure 5f**)

A recent study of mouse Atp2b2 revealed multiple conformations bound to phosphatidylinositol 4,5-bisphosphate (PIP_**2**_), implicating it as an allosteric regulator of the ATP-driven P-type ion transport^88^. To test robustness across conformational states, we mapped human variants onto homologous residues in mouse Atp2b2 structures and applied 3DNT independently to each state of the Post-Albers cycle. Enriched neighborhoods remained near the Ca^2+^ binding site and the adjacent PIP_2_ interface in all states (Figure S8).

### Neuropsychiatric-associated ATP2B2 variants disrupt ATP- and PIP_2_-driven conformational control of the functional cycle

Having established consistency between different structural models, we sought to interpret variants disrupting key ligand binding sites and interdomain interfaces before examining variants affecting PIP_2_-mediated regulation.

The conserved aspartate D733 is a putative mediator of ATP-dependent domain rearrangements. ATP binding drives a concerted transition from an open, flexible configuration of the cytoplasmic N, P, and A domains to a compact, closed E1-ATP conformation primed for autophosphorylation, and mechanically coupled to Ca^2+^ binding and occlusion in the membrane. In the ATP-bound state, D733 localizes to the N-P-A domain interface, stabilizing the closed conformation. The schizophrenia-associated D733N variant substitutes the negatively charged aspartate with neutral asparagine while conserving side chain volume, selectively disrupting electrostatic interactions at this interdomain interface. Electrostatic surface analysis indicated a sizable change in electrostatic potential within a 7 Å radius relative to wild-type (Figure S7a).

We next examined variants lying near the PIP_2_ interaction interface, where disruption of PIP_2_-mediated regulation may underlie their effects. PIP_2_ stabilizes the Ca^2+^-bound E1 state through direct interaction with N866, whose engagement with PIP_2_’s fatty acid ester bond prevents interference with the adjacent binding site—acting as a molecular latch that couples lipid binding to pump state. V885I, a variant identified in a schizophrenia case, lies immediately adjacent to N866 (**Figure S7c**), suggesting that it disrupts this latch and impairs PIP_2_-mediated regulation of pump cycling.

A second schizophrenia case variant, A400E, also maps near the PIP_2_ molecule (**Figure S7c**). A400 contributes to formation of the PIP_2_ binding pocket as part of a hydrophobic wall stabilizing the lipid acyl chains. Missense3D analysis of both the cryo-EM and AlphaFold3-predicted structures predicted that A400E (replacing a small alanine side chain with a negatively charged glutamate) enlarged the local cavity by about 174 Å^3^, potentially destabilizing PIP_2_ accommodation.

Nearby, a *de novo* variant, S922L, replaces a small, polar serine with a hydrophobic leucine in a transmembrane segment positioned adjacent to the putative counter-proton release pathway. Missense variants within this pathway (S877F, associated with hereditary deafness^89^) severely impair Ca^2+^ transport by disrupting the proton movements that are coupled to Ca^2+^ extrusion and required to reset the ion-binding site during the Post-Albers cycle^82^. Missense3D analysis of the AlphaFold3-predicted structure indicated that S922L expands the local cavity volume by 378.0 Å^3^, consistent with disruption of the tightly packed transmembrane architecture required for Ca^2+^ extrusion. Interestingly, SERCA pump mutations associated with Darier’s Disease also affect a conserved lipid binding site, and the same site is affected as well by some of the mutations in the neuron-specific ATP1A3 sodium-potassium pump that are associated with alternating hemiplegia of childhood^90^.

Together, these observations suggest that a subset of missense variants in ATP2B2 may impair pump function not only by directly perturbing Ca^2+^ binding or catalytic domains, but also by disrupting lipid-mediated regulatory mechanisms essential for transport.

### Functional testing of missense variants

To evaluate variants at Ca^2+^-binding and ATP:Mg^2+^ sites, we selected substitutions from our 3D neighborhood analysis in closest proximity to each ligand. Given that ATP hydrolysis is the central catalytic function of ATP2B2, we assessed whether these variants impaired enzymatic activity using a colorimetric ATPase assay that quantifies inorganic phosphate release in the PMCA2z/a isoform (Figure 6a; Methods).

**Figure 6:**
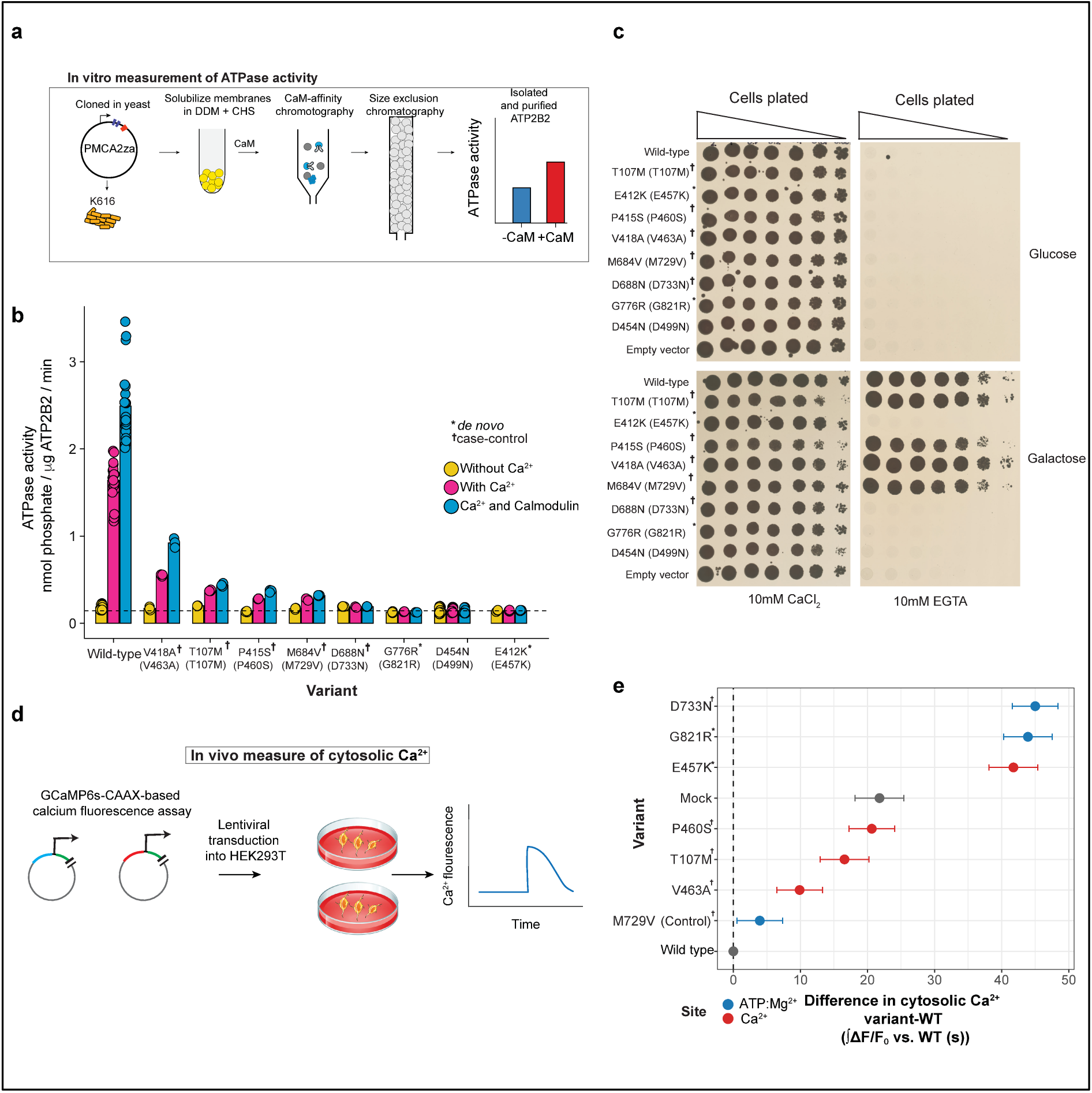
Functional characterization of ATP2B2 missense variants prioritized by structural analysis. A. Overview of the measurement of ATPase activity of schizophrenia, NDD, and ASD missense variants in ATP2B2. An isoform of ATP2B2, PMCA2z/a, was expressed in yeast, membranes were solubilized in DDM and CHS, and the protein was purified by calmodulin affinity chromatography followed by size exclusion chromatography. (see Methods: In Vitro Measurement of ATPase Activity). B. ATPase activity is shown for wild-type and mutant ATP2B2 proteins under three conditions: w/o Ca^2+^ (yellow), Ca^2+^ (pink), and Ca^2+^/CaM (blue). Variants located closest to the Ca^2+^ or ATP:Mg coordination sites show the most pronounced functional deficits. E457K, D733N, and G821R exhibit near complete loss of ATPase activity across conditions, comparable to the catalytically inactive control D454N, consistent with disruption of core catalytic and ion binding elements. In contrast, P460S and V463A retain partial Ca^2+^- and Ca^2+^/CaM-stimulated ATPase activity, indicating preserved but reduced catalytic capacity. The control variant M684V similarly maintains measurable stimulated activity. Bars represent mean activity and points represent individual measurements. The dashed horizontal line indicates the average activity of the dead variant values. Variants marked with † are derived from schizophrenia cases, and those marked with * are derived from NDD or ASD cases. M684V serves as the control variant. C. Yeast complementation assay assessing the functional impact of ATP2B2 missense variants. Yeast K616 expressing wild-type or mutant ATP2B2 were plated on glucose or galactose media in the presence of 10 mM CaCl_2_ or 10 mM EGTA. Growth on galactose reflects ATP2B2-dependent calcium handling, while loss of growth indicates impaired pump function. Variants retaining ATPase activity, including T107M, P460S (P415S), V463A (V418A), and M729V (M684V), supported growth under Ca^2+^-limiting conditions, whereas catalytically inactive variants E457K (E412K), D733N (D688N), and G821R (G776R) failed to complement growth and phenocopied the empty vector control. D. Overview of ATP2B2 missense variant effects on cytosolic calcium responses in HEK293T cells. Synthesized DNA constructs containing ATP2B2 alleles were inserted into a lentiviral vector and transduced into HEK293T cells engineered to express Kir2.3 channels and the membrane-tethered genetically encoded calcium sensor GCaMP6s-CAAX. Following transduction, cells were hyperpolarized and intracellular calcium levels were then quantified using a high-throughput fluorometric imaging plate reader (see Methods: Testing ATP2B2 variants in HEK cells). E. Similar to 6B, variants located proximal to the Ca^2+^ or ATP:Mg^2+^ coordination sites resulted in the highest amount of cytosolic calcium (E457K, D733N, and G821R). Dots indicate model estimated mean differences in calcium response relative to wild-type from a linear mixed effects model, with error bars showing 95% confidence intervals.

Variants most proximal to the catalytic core (E412K, D688N, and G776R in the PMCA2z/a isoform; E457K, D733N, and G821R) exhibited complete loss of ATPase activity under Ca^2+^-saturating conditions, both in the absence and presence of calmodulin, phenocopying a catalytically inactive mutant (Figure 6b). In contrast, P460S (P415S) and V463A (V418A), located more distally, retained measurable activity, as did M729V (M684V), a variant observed exclusively in unaffected controls. Nevertheless, all variants showed substantial reductions relative to wild type, with residual activity decreased by approximately 5–10-fold and several approaching baseline.

We next assessed Ca^2+^ transport capacity using a yeast complementation assay in the Ca^2+^ transport-deficient strain K616, which requires expression of a functional Ca^2+^ pump to support growth under Ca^2+^-limiting conditions (Methods). Variants disrupting the Ca^2+^-binding site or catalytic function failed to rescue growth under EGTA-imposed Ca^2+^ depletion (Figure 6c). In contrast, variants that retained partial ATPase activity supported growth, broadly consistent with the enzymatic defects observed in vitro.

Because cellular Ca^2+^ homeostasis is regulated by coordinated activity across multiple pathways, we next evaluated variant effects in a physiologically relevant cellular context using a genetically engineered HEK293T-GCaMP6s-CAAX system optimized for submembrane Ca^2+^ monitoring^91^. PMCA2z/a variants were delivered by lentiviral transduction using PGK-driven constructs containing an upstream mCherry-P2A cassette for expression-matched sorting (Figure 6d). Western blot analysis confirmed similar expression levels across all transfected variants (Figure S10b). Cells were depolarized with KCl at EC_10_, and variant effects on cytosolic Ca^2+^ responses were quantified using a linear mixed-effects model of fluorescence AUC relative to wild-type (Methods).

Consistent with their structural location, E457K and G821R, both *de novo* variants, together with the case schizophrenia variant D733N, produced the largest increases in cytosolic Ca^2+^ (ΔAUC ~40–45 relative to wild type), indicating severe impairment of Ca^2+^ extrusion. Variants positioned more distally exhibited more modest increases (ΔAUC ~10–20), while the control variant most closely resembled wild-type ATP2B2 (Figure 6e). Together, these results indicate that missense variants associated with key neuropsychiatric disorders impair ATP2B2 through two independent mechanisms—disruptions to Ca^2+^ coordination or ATP-dependent catalysis—with functional severity scaling with proximity to ligand-binding sites, ultimately resulting in defective calcium extrusion.

## Discussion

We found that genes that encode the proteins that manage Ca^2+^ fluxes at synapses were disproportionately enriched for genetic variation that affects risk neuropsychiatric disorders, an enrichment that exceeded not only that for other ion channel classes, but also the canonically implicated gene sets including the targets of FMRP and CHD8^62,92–95^(Figure 2c). These findings suggest that precise control of Ca^2+^ fluxes and their duration and magnitude is important in protection from these disorders, and that this protection is sensitive to a wide variety of common and rare genetic perturbations.

We then sought to better understand a specific part of this signal, through a structure-informed genetic analysis of PMCA2 (encoded by *ATP2B2*), a transporter that extrudes Ca^2+^ from synapses. We found that missense variants in ATP2B2 segregated into two spatially distinct regions of the protein: (1) the Ca^2+^ transport substructure, which includes the canonical Ca^2+^-binding site and residues involved in PIP_2_-dependent regulation of pump activity, and (2) the ATP:Mg^2+^-binding pocket required for catalytic cycling. This spatial, structural concentration of the genetic signal from missense variants supports two kinds of dysfunction, in which variants impair either Ca^2+^ transport itself or the ATP hydrolysis that supports this transport energetically. These patterns were consistent across AlphaFold3 models and a well-determined cryo-EM structure and were further supported by electrostatic and ligand interaction analyses. Multiple functional assays showed that variants from both regions reduce ATPase activity and impair Ca^2+^ extrusion, resulting in sustained elevation of cytosolic Ca^2+^ (Figure 6).

PMCAs create and restore the steep calcium gradient that exists across synaptic membranes, keeping intracellular Ca^2+^ at ~100 nM despite much higher extracellular levels^57,58,67,96,97^. This gradient enables Ca^2+^ channels (when opened) to elicit rapid Ca^2+^ transients, that in turn instruct neurotransmitter release, dendritic integration, synaptic plasticity, and activity-dependent gene expression^98–101^. More specifically, the genetic findings implicating both efflux (plasma membrane Ca^2+^ transporters) and influx (voltage-gated Ca^2+^ channels) indicate that precise regulation of Ca^2+^ dynamics – the magnitude, duration and perhaps the shape of these Ca^2+^ transients – is critical.

Disruption of Ca^2+^ dynamics would broadly affect neuronal signaling, producing responses of inappropriate magnitude or timing. Several effectors of Ca^2+^-dependent transcription, including CREB and members of the SWI/SNF complex, themselves carry neuropsychiatric genetic signal^34,38,72,94^. ATP2B2 is further modulated by membrane PIP_2_, which phospholipase C cleaves to IP_3_ during receptor-mediated signaling^102,103^, periods of heightened synaptic activity could thus deplete PIP_2_ and attenuate pump activation.

Beyond rare patients with ATP2B2-disabling mutations, how broadly might people with schizophrenia exhibit reduced ATP2B2 expression? We tested this in two distinct, case-control datasets (one transcriptomic^23^ and one proteomic^104^), from the dorsolateral prefrontal cortex (DLPFC; Brodmann area 46). In single-nucleus RNA-seq from 191 donors (94 with schizophrenia and 97 controls)^23^. ATP2B2 was significantly reduced specifically in glutamatergic neurons, (**Figure S12a**). In a separate proteomic dataset of synaptic fractions from 69 donors (34 with schizophrenia and 35 controls)^104^, ATP2B2 was likewise reduced in cases (**Figure S12b**). Similar downregulation was observed for other postsynaptic density genes implicated in psychiatric risk (*GRIA3, SHANK2, GSK3B*; Figure (**Figure S12d**), which are among the large set of genes regulated by the Synaptic Neuron-Astrocyte Program (SNAP^23^). These findings support a pathogenic model in which either rare missense variants that perturb Ca^2+^ efflux, or common biological changes (such as a decline in SNAP affecting expression of ATP2B2 and other synaptic components), contribute to schizophrenia etiology.

Methodologically, we introduced a structure-informed framework for analyzing and prioritizing among missense variants, recognizing the biology on which they converge and strongly informing experimental follow-up. This approach is not limited to any prior functional hypothesis and does not depend on predefined functional annotations of protein domains but rather uses unbiased identification of functionally critical regions directly from protein structure itself. Rather than subjecting all candidate variants to labor-intensive functional assays, it prioritizes those variants with the highest likelihood of pathogenicity. Applied to ATP2B2, prioritized variants showed significantly higher AlphaMissense pathogenicity scores than variants outside these regions, both across all variants (Wilcoxon P = 8.98 × 10^−9^) and when restricted to cases alone (Wilcoxon P = 1.1 × 10^−3^). Critically, this reduced the number of variants requiring experimental validation from 336 to 54—a 6.2-fold reduction. By comparison, applying a uniform AlphaMissense threshold (≥0.564^17^) would retain ~46% of sites, highlighting the substantially greater specificity of structure-informed prioritization, likely because AlphaMissense primarily captures evolutionary constraint rather than phenotype-specific functional perturbation. This demonstrates that a protein-structure-guided approach can reduce exhaustive variant-by-variant experimental testing without sacrificing biological relevance. Given remarkable recent progress in the ability to predict protein structures computationally, we hope that this approach can be broadly helpful in connecting human genetic data sets to specific protein functions.

### Limitations

Our work also has limitations. First, current single-cell transcriptomic and human genetic datasets capture only part of the genetic complexity of polygenic neuropsychiatric traits, and our focus on neurons does not preclude contributions from other brain cell types. Second, the 3D neighborhood test depends on structural model accuracy and has limited power in flexible or poorly resolved regions (e.g., PDZ-binding domains). The test identifies regions where case variants are disproportionately concentrated rather than isolating individual causal variants and should therefore be interpreted as variant prioritization rather than definitive evidence of pathogenicity. Larger sequencing studies will improve power, and variants observed only in controls are not necessarily benign — they may be less deleterious or modulated by polygenic background and environmental risk. Finally, yeast and HEK293 systems lack neuronal signaling architecture; dendritic spines, postsynaptic density organization, activity-driven calcium microdomains, neuron-specific cofactors and lipid environments — so endogenous ATP2B2 regulation, localization, and coupling to native calcium channels are only partially captured.

## Figures

**S1.**
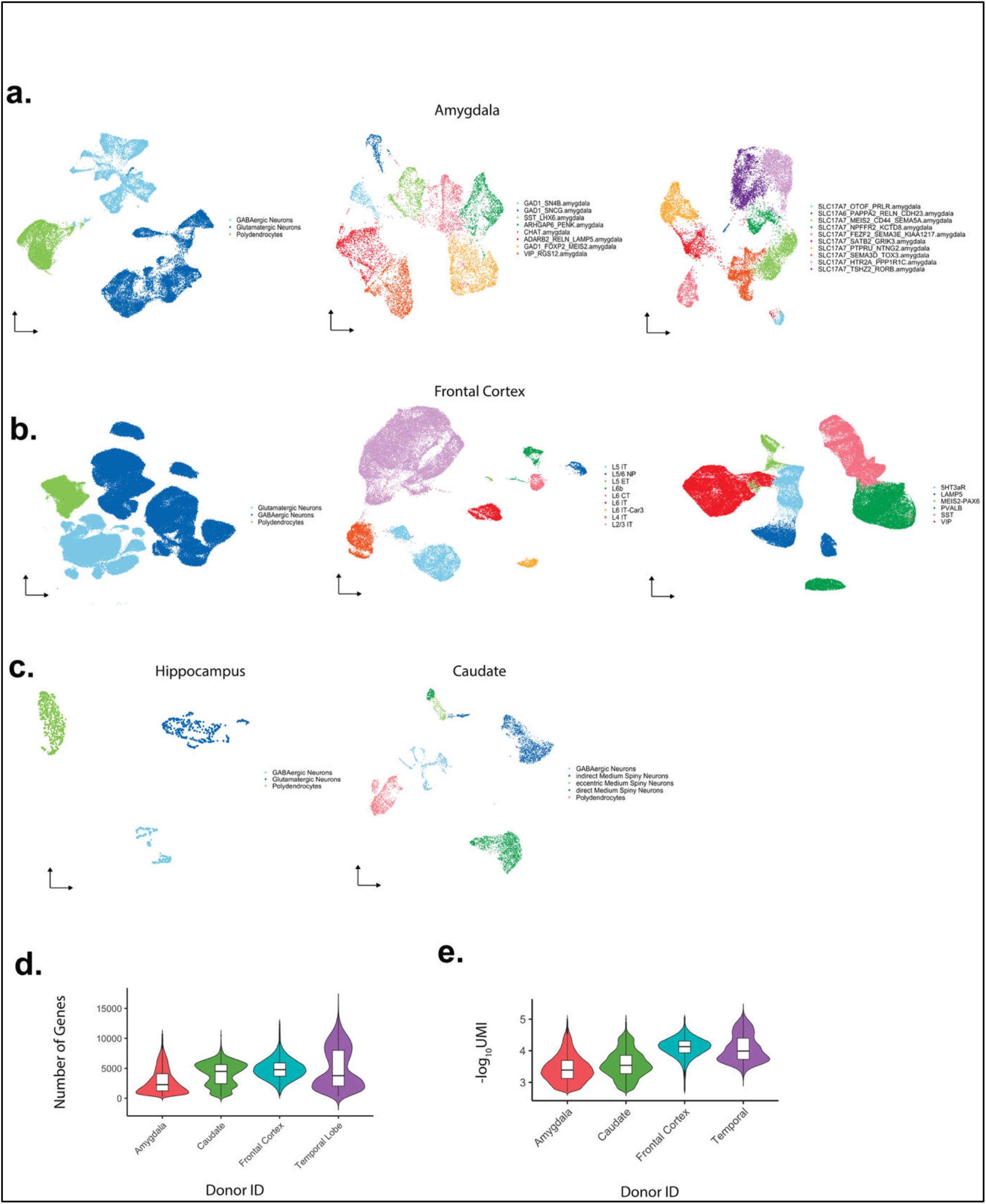
Brain single cell datasets used in this study. A. A-D UMAP embeddings of scRNA-seq profiles colored by cell type annotations. Amygdala (A) and frontal cortex (B), neuronal subtypes are included. B. Number of genes (d) and UMIs (unique molecular indicators) grouped by region.

**S2.**
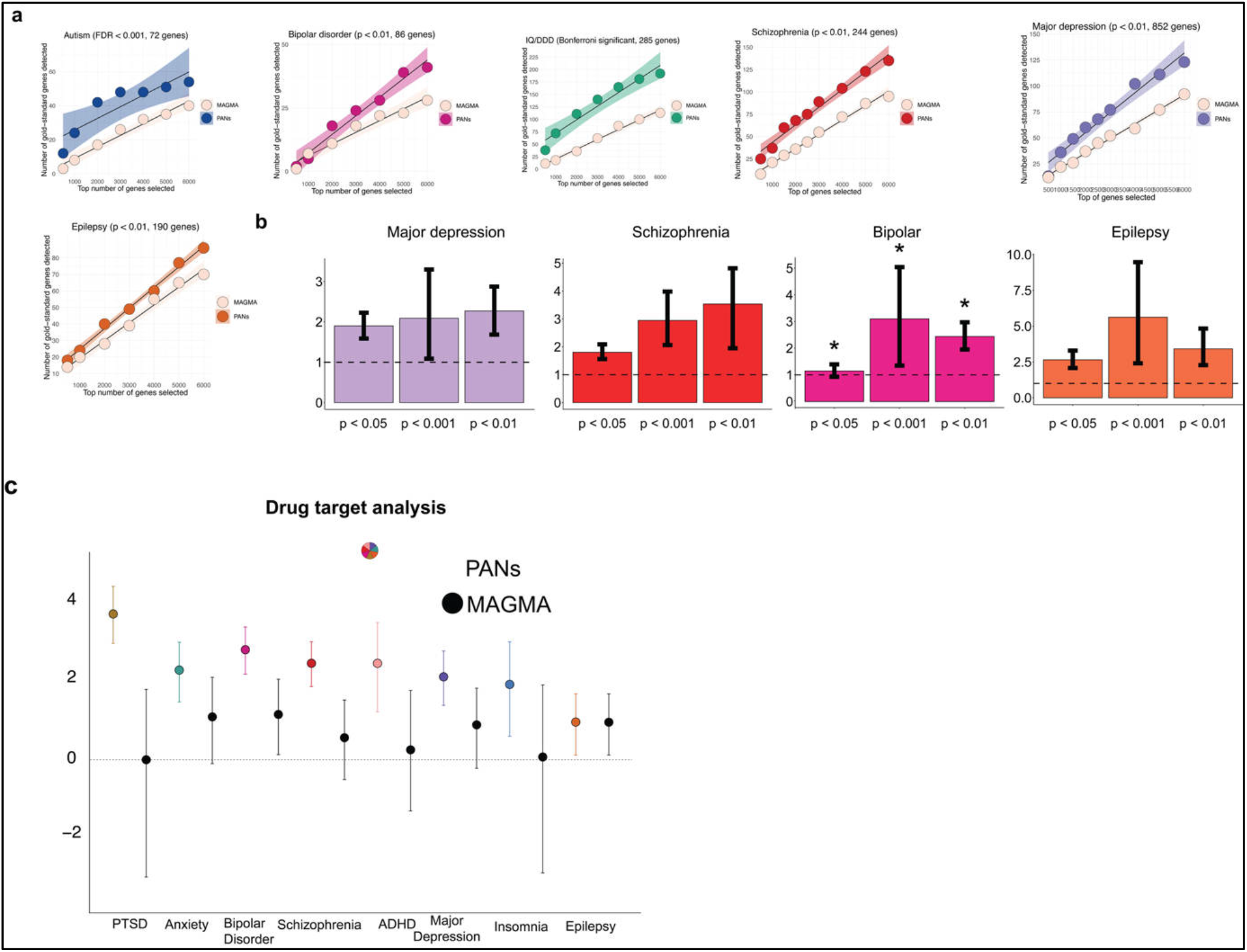
Genetic analyses of prioritized genes in exome data. A. For neuropsychiatric traits with available exome sequencing data, our pairwise prioritization approach identifies a greater number of genes across independent exome studies compared to MAGMA. The x axis represents the top N genes from each method’s ranked list. For neurodevelopmental disorder and autism, significance thresholds were defined according to the original criteria reported in the respective studies. For schizophrenia, bipolar disorder, major depressive disorder, and epilepsy, a P value threshold of 0.01 was applied. For MAGMA, we included protein coding genes with Z score greater than 0 and P value. B. Fold enrichment of prioritized gene scores in exome sequencing data for neuropsychiatric traits across multiple P value thresholds is shown, analogous to Figure 2a. Error bars represent 95 percent confidence intervals derived from bootstrapping PAN scores for significant versus non-significant genes within each phenotype, as described in the Supplementary Methods. Statistical significance is indicated by asterisks corresponding to Wilcoxon rank sum tests comparing PAN scores between significant and non-significant genes at each sequencing threshold. C. Reported −log_10_ p-values were obtained by applying Fisher’s method on the two-sided P values of both protein truncating and damaging missense variants to constrained drug target sets from schizophrenia, and just damaging missense variants for bipolar disorder (Numerical results are in Table S3). The dot represents the OR and the bar represents the 95% CI of the point estimates.

**S3.**
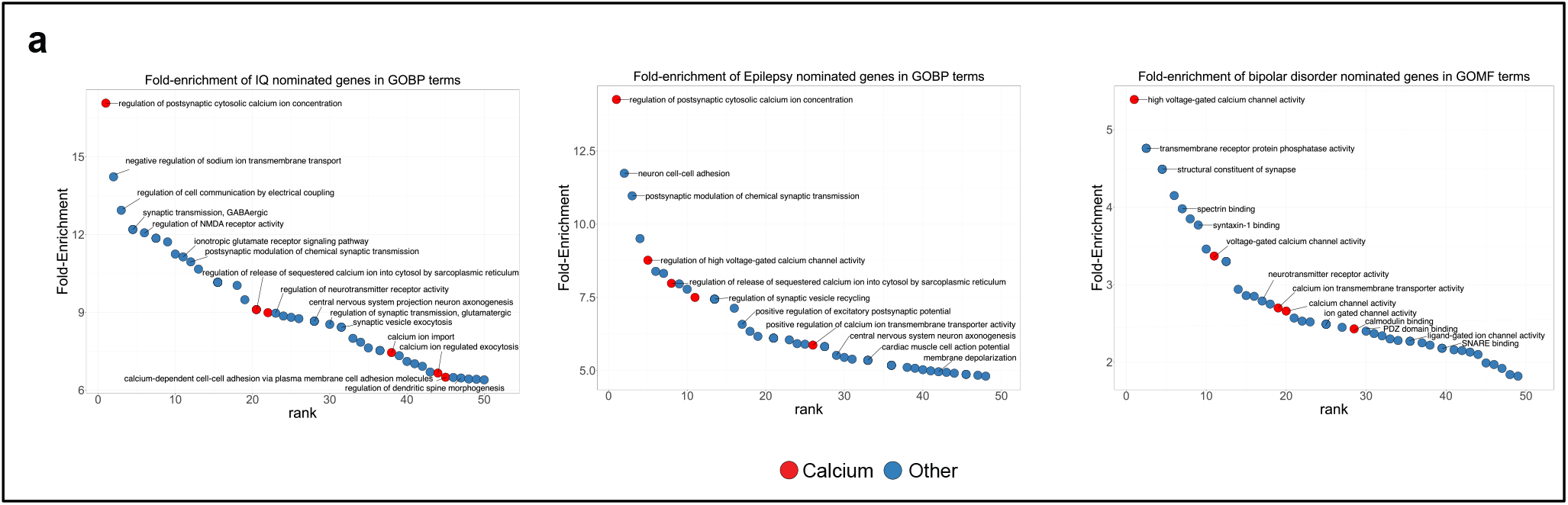
Gene ontology analysis of other brain traits. A. Fold-enrichment of nominated genes in Gene Ontology terms in IQ, Bipolar Disorder, and Epilepsy (corresponding to Figure 2a). Terms related to calcium signaling and homeostasis (red) are notably enriched. In all panels, terms not related to calcium are shown in blue.

**S4.**
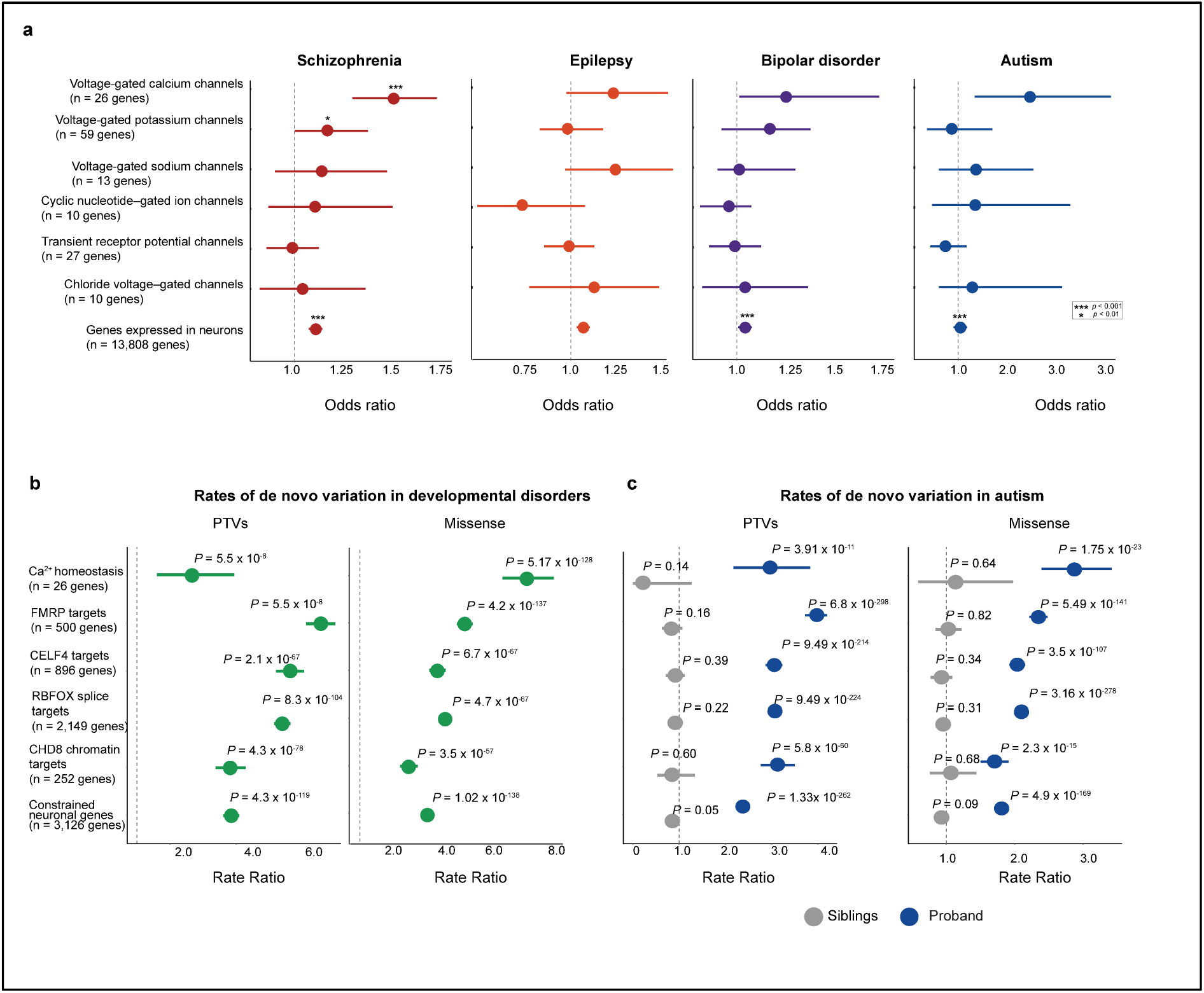
Genetic enrichment of voltage-gated ion channels and Ca^2+^ flux pathway genes across neuropsychiatric and neurodevelopmental disorders. A. Case-control enrichment and excess case rare-variant burden in Ca^2+^, K^+^ and Na voltage gated channel genes (corresponding to Figure 2e) in epilepsy, autism and bipolar disorder. The dot represents the OR and the bar represents the 95% CI of the point estimates. We used neuronal genes as a background, defined as genes that are expressed in at least 5% of any one neuronal cluster from single cell RNA-seq data. Gene lists obtained from the HUGO gene nomenclature committee (HGNC), unlike the analyses in Figure 2, we used the entire gene set from the HGNC, and did not filter on constraint. B. Rates of de novo protein-truncating (left) and missense (right) variants observed in developmental disorders. Compared to other gene sets specific to nervous system biology, constrained members of the Ca flux pathway genes exhibit marked enrichment for missense variation. Enrichment for protein-truncating variants in Ca^2+^ flux pathway genes is also significant. C. Data are from the 2025 (unpublished) Autism Sequencing Consortium (ASC) analysis showing *de novo* rates in probands (blue) versus neurotypical siblings (grey). Compared to other gene sets relevant to nervous system function, constrained members of the Ca^2+^ flux pathway gene set exhibit enrichment for missense variation. Enrichment for protein-truncating variants in Ca^2+^ flux pathway genes is also observed, though more modest relative to other gene sets.

**S5.**
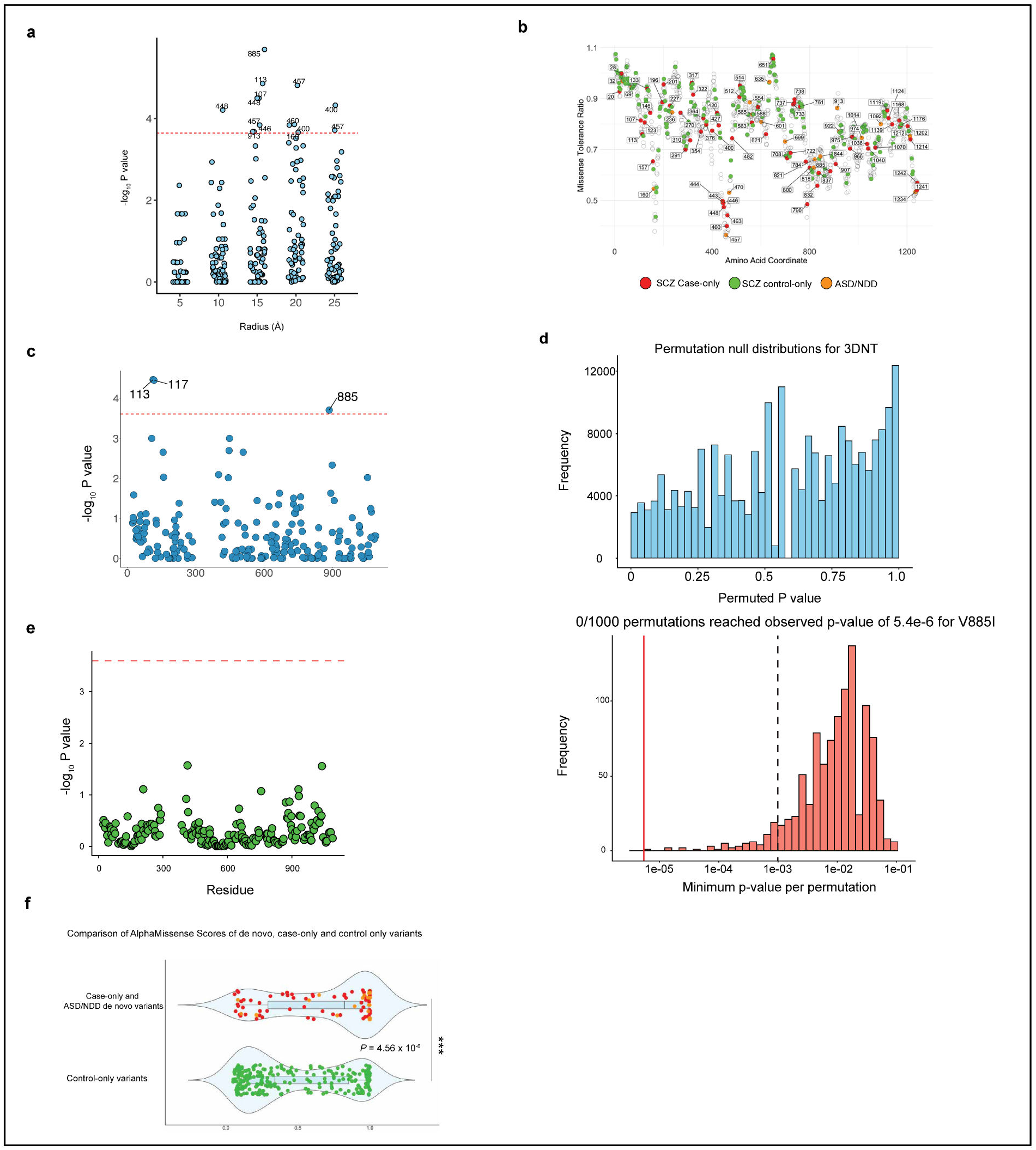
Residue analysis of ATP2B2. A. Sensitivity of the 3D neighborhood analysis to neighborhood radius B. Missense tolerance ratio (MTR) ^13,105^ track of ATP2B2 from the Regeneron Genetics Center Million Exome dataset ^13^. Residues with highly damaging variants associated with schizophrenia case-only, ASD or NDD studies are colored. Glu457 is a residue located within one of the most regionally constrained regions of the gene. The data is taken from the Regeneron Genetics Center (RGC) browser. C. P values from the chi square test of P-values from the per-residue neighborhood burden enrichment analysis, subsetted just to case-only or control-only variants from the largest schizophrenia exome study. The red line represents the Bonferroni significance threshold, with labeled residues indicating those reaching statistical significance. D. Case–control labels were shuffled 1,000 times, preserving the observed set of variant residue positions and overall case/control counts, and the 3D neighborhood test was re-run at every qualifying residue. Top: Pooled null: Fisher’s exact one-sided p-values across all 219 residues × 1,000 permutations (n = 219,000). The distribution is non-uniform because tests share carriers and neighborhoods, but no permuted p-value reached the observed V885I value (P_obs = 5.4 × 10^−6^ vertical line). Botton: Family-wise null: minimum p-value across the 219 residues for each of the 1,000 permutations. The 5th-percentile FWER threshold was 8.94 × 10^−10^ and 0/1,000 permutations produced a minimum p-value ≤ P_obs, giving FWER-corrected P_perm < 10^−3^. E. 3D neighborhood analysis of synonymous variants from schizophrenia exome sequencing data. No residues exhibit Bonferroni-significant clustering of synonymous case variants. The red line represents the Bonferroni significance threshold, with labeled residues indicating those reaching statistical significance F. Comparisons of AlphaMissense pathogenicity scores of the non-filtered case-only schizophrenia, ASD and NDD-associated variants to control-only variants. The P value is computed from a one-sided Wilcoxon test.

**S6.**
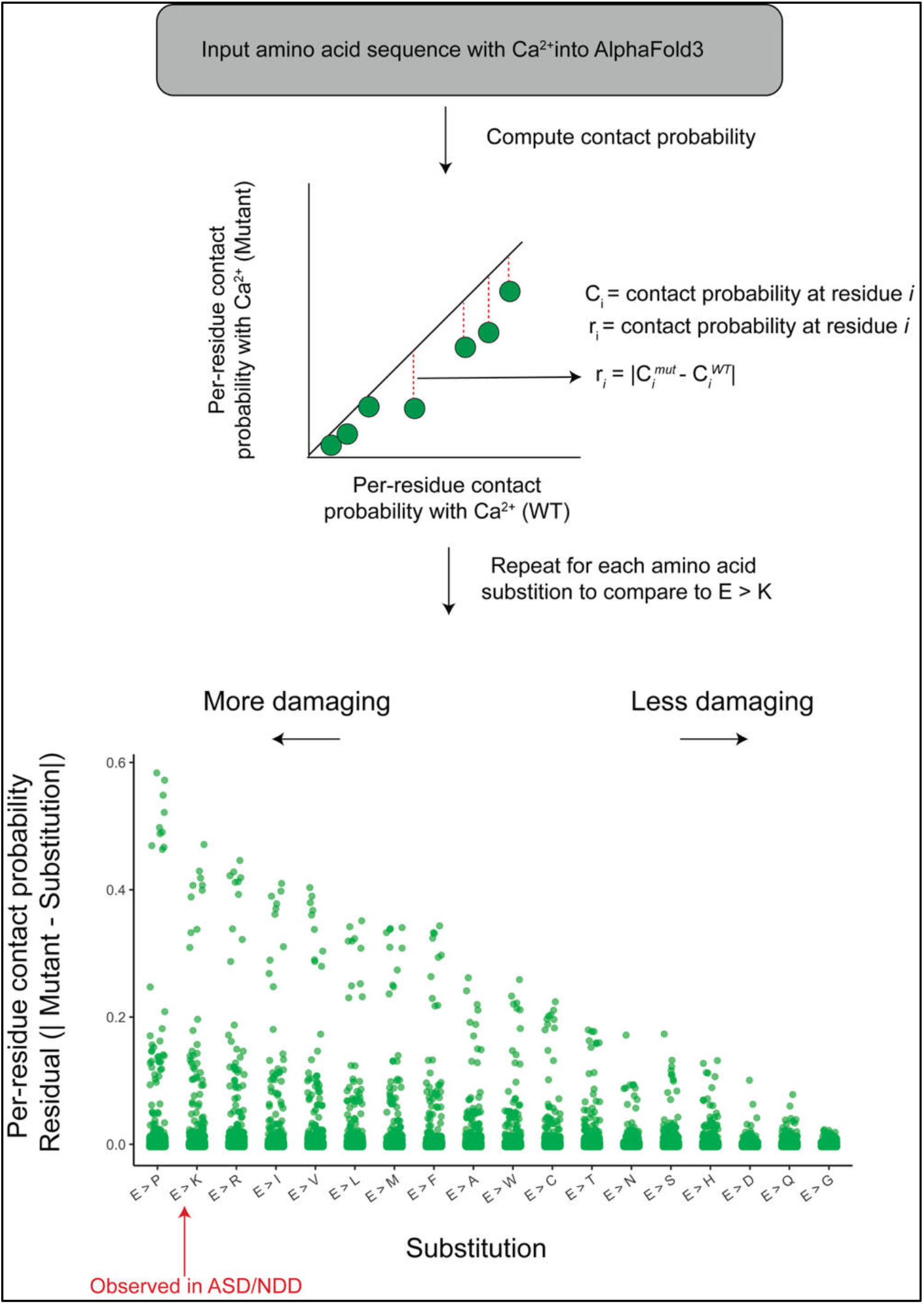
Effect of All Possible E457 Mutations on Residue-Level Calcium Contact Probability in ATP2B2. A. Top. Workflow illustrating how AlphaFold3 was used to model mutant structures of ATP2B2 with bound Ca^2+^, compute per-residue calcium contact probabilities, and quantify disruption relative to wild type. Residual contact disruption was calculated for each residue. Bottom: Distribution of per-residue residuals for all substitutions at glutamic acid 457 (E457), sorted by the maximum disruption observed per mutation. Mutations are ordered from most to least structurally disruptive to calcium contacts. Substitutions that differ substantially from glutamic acid in charge, size, or hydropathy—such as proline and arginine—were among the most disruptive. In contrast, substitutions to chemically similar amino acids, such as glycine, glutamine, and asparagine, caused minimal disruption. The E>K mutation, which was seen in probands with autism and neurodevelopmental disorders (ASD/NDD), is among the most disruptive.

**S7.**
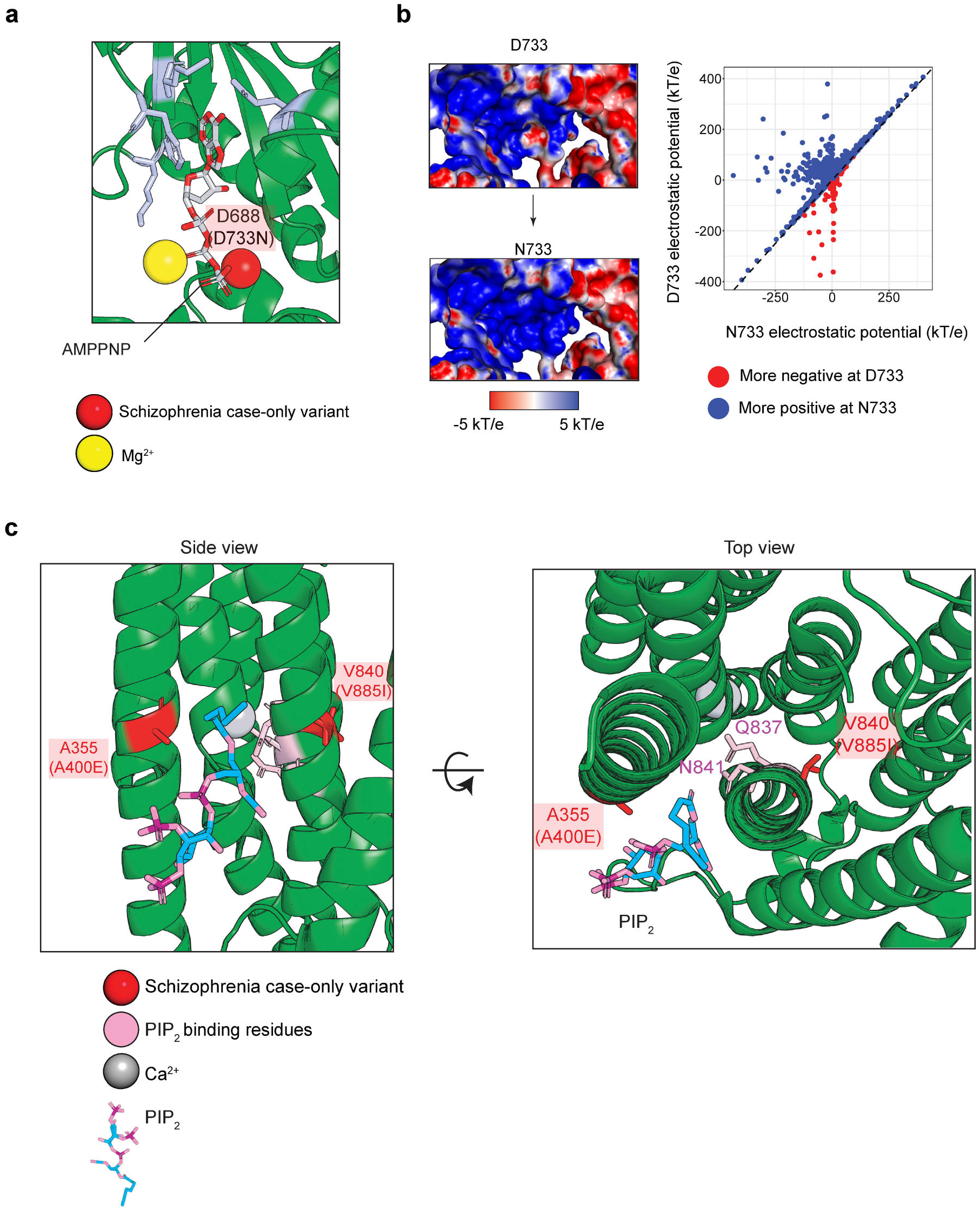
Structural and electrostatic consequences of schizophrenia case-only variants in ATP2B2. A. Nucleotide binding occurs at the interface of the three cytoplasmic domains following the E1 to E1-ATP transition. Shown is the structure of mouse ATP2B2 (PDB: 9GSF), in the E1-ATP state, used here for illustration as this conformation was not captured in our human PMCA2z/a cryo-EM structure. The non-hydrolyzable ATP analog AMPPNP is bound at the N, P, and A domain interface and coordinated by Mg^2+^. Residues E525, F557, K562, and M564 (mouse numbering) are highlighted in blue to indicate direct nucleotide contacts. D688 (mouse), corresponding to D733 in human PMCA2z/a, is the site of the schizophrenia-associated D733N variant. This residue lies adjacent to the nucleotide binding interface, suggesting that the D733N substitution may alter local electrostatics or N-P domain coupling during ATP binding, thereby perturbing catalytic activity. B. Electrostatic surface comparison of the D733 and N733 states, with a scatter plot quantifying residue-level electrostatic potential differences, highlighting regions higher in D733 or N733.Surface representation of ATP2B2 is colored by electrostatic Coulomb potential, ranging from –5 kcal (mol e)^−1^ in red to +5 kcal (mol e)^−1^ in blue. C. Membrane-proximal region showing schizophrenia case-only variants A355 (corresponding to A400E in human PMCA2z/a) and V840 (corresponding to V885I in human PMCA2z/a) positioned directly adjacent to PIP_2_ binding residues and a bound calcium ion, suggesting these variants may disrupt lipid binding and calcium coordination at this regulatory interface. Residue numbering follows mouse ATP2B2 (PDB: 9GSI).

**S8.**
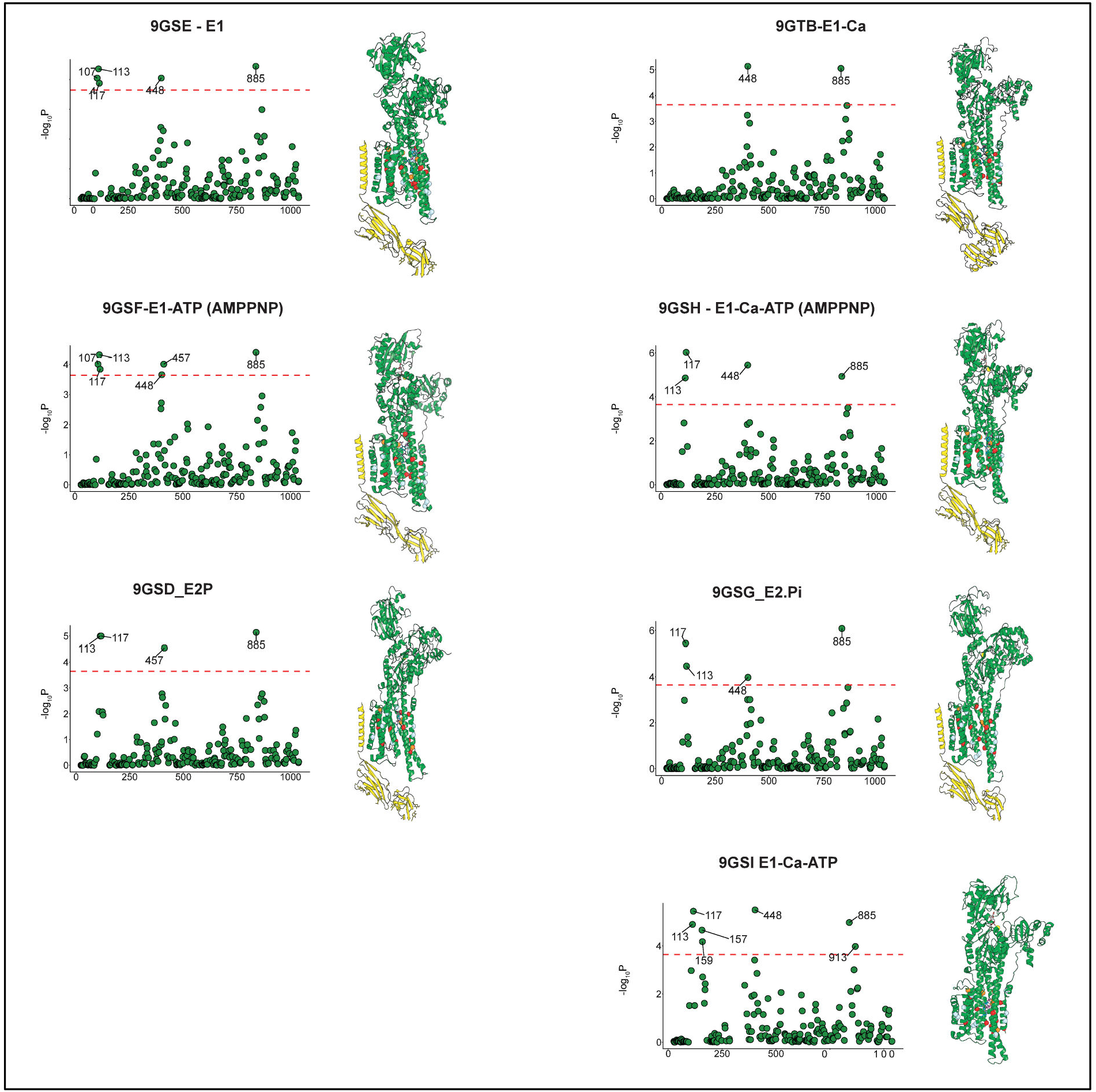
3D neighborhood enrichment of missense variants in mouse PMCA2 is conserved across Post-Albers cycle conformations and matches AlphaFold predictions. A. Cryo-EM structures of mouse PMCA2 spanning the Post-Albers transport cycle (E1, E1-Ca, E1-ATP, E1-Ca-ATP, E2P, and E2-Pi; PDB IDs indicated) were used to perform a 15 A 3D neighborhood scan for enrichment of missense variants. Scatter plots show residue position (mouse/PDB numbering) versus −log10(p) from one-sided Fisher’s exact test, with the red dashed line denoting the Bonferroni-corrected significance threshold. Significant residues are labeled by position in human PMCA2 (Q01814). Corresponding cryo-EM models are shown alongside each plot.

**S9.**
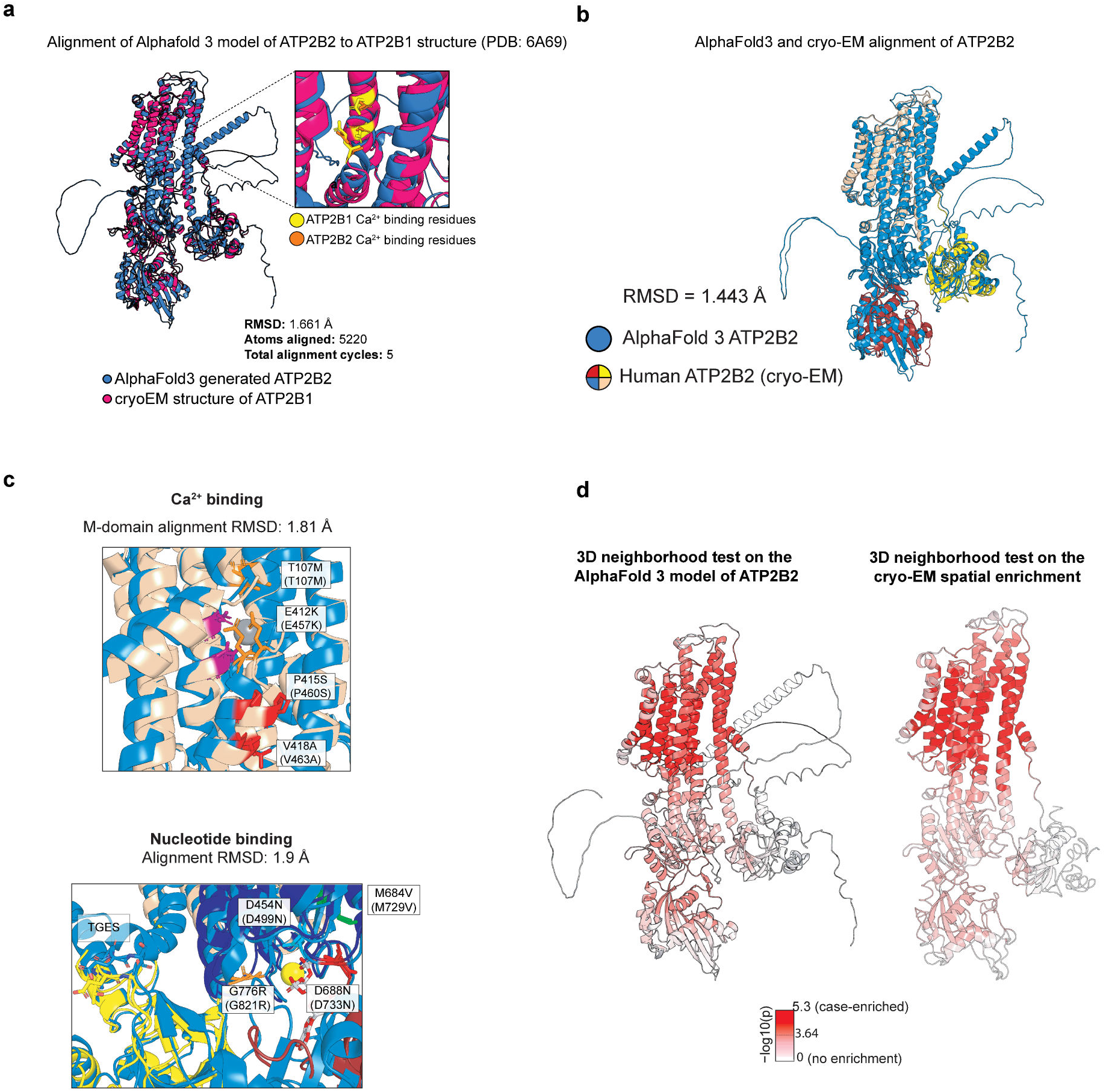
Cryo-EM structure determination and validation of human ATP2B2 (PMCA2z/a). A. Structural alignment of the ATP2B2 AlphaFold3 model with the ATP2B1 cryo-EM was performed to identify the Ca^2+^ binding site. AlphaFold3 and reference structures were aligned on Ca atoms, restricting analysis to residues with PLDDT greater than 70 stored in the B factor field. B. Structural alignment of ATP2B2 AlphaFold3 model with cryo-EM structure. AlphaFold3 and reference structures were aligned on Cα atoms, restricting analysis to residues with pLDDT greater than 70 stored in the B factor field. C. Structural alignment of the M domain between AlphaFold and cryo-EM structures showing high concordance (M-domain RMSD = 1.81 Å). Mutated residues around the nucleotide binding site are in stick representation. D. Spatial enrichment map in both the cry-EM model and the AlphaFold 3 model of full-length human ATP2B2. Color scale capped at –log_10_(p) = 5.3.

**S10.**
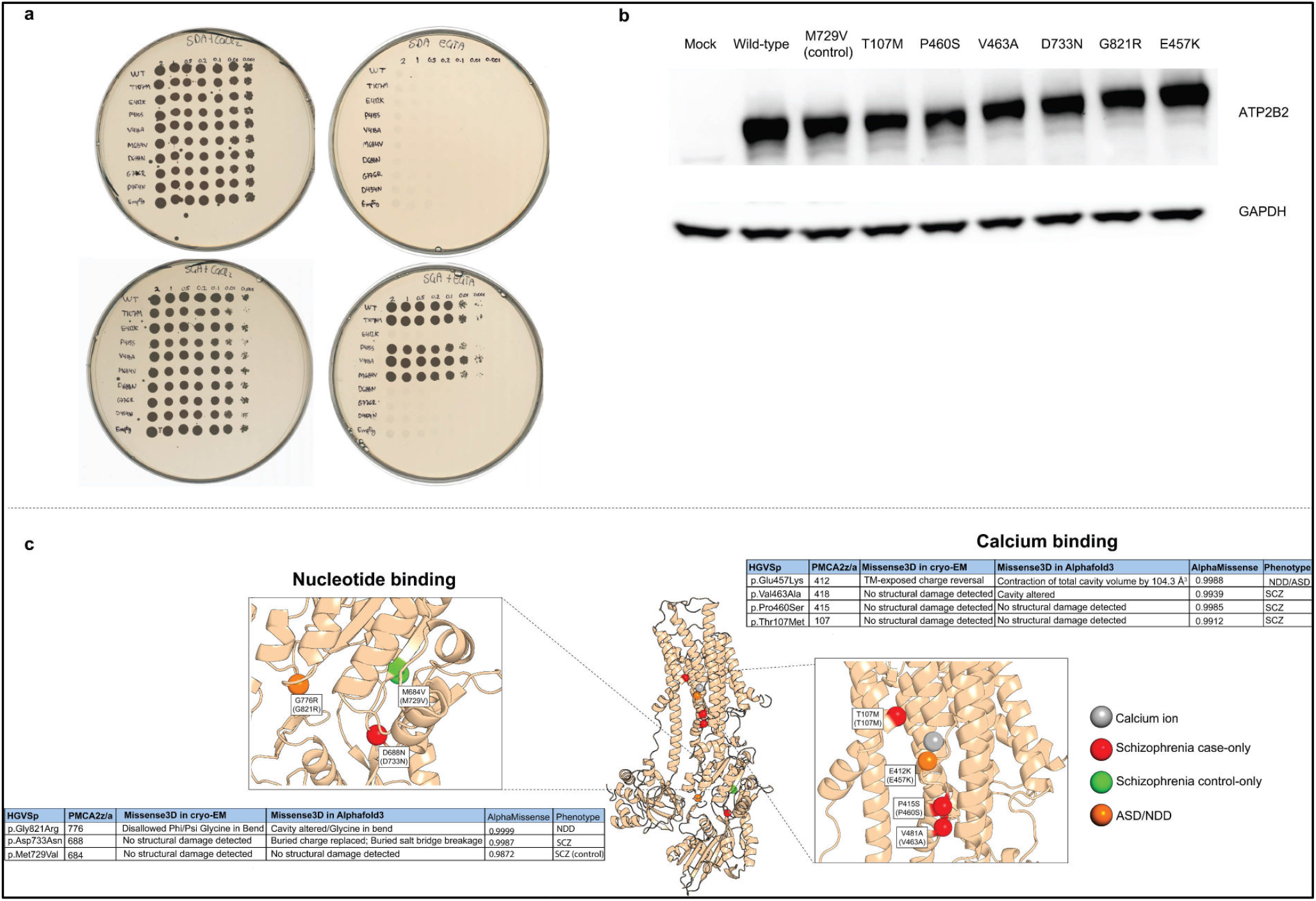
Yeast complementation raw data and protein expression controls for ATP2B2 variants. A. Unprocessed yeast complementation plate images corresponding to Figure 3. Serial dilutions of yeast expressing wild-type or mutant ATP2B2 were plated under repressing (glucose) or inducing (galactose) conditions in the presence of CaCl_2_ or EGTA. D454N serves as a catalytically inactive negative control. B. Western blot analysis of ATP2B2 expression in cell lysates. Immunoblotting for ATP2B2 shows comparable protein expression across wild-type and variant constructs, with GAPDH serving as a loading control. These data indicate that functional differences observed for specific variants are not explained by gross differences in protein abundance. C. Structural mapping of variants selected for biochemical and cellular assays onto the cryo-EM model, showing their spatial positions relative to key functional domains (related to figure 6). Missense3D was applied to the cryo-EM structure, which contains one bound Ca^2+^, and to the AlphaFold3 models containing either Ca^2+^ or ATP·Mg^2+^.

**S11.**
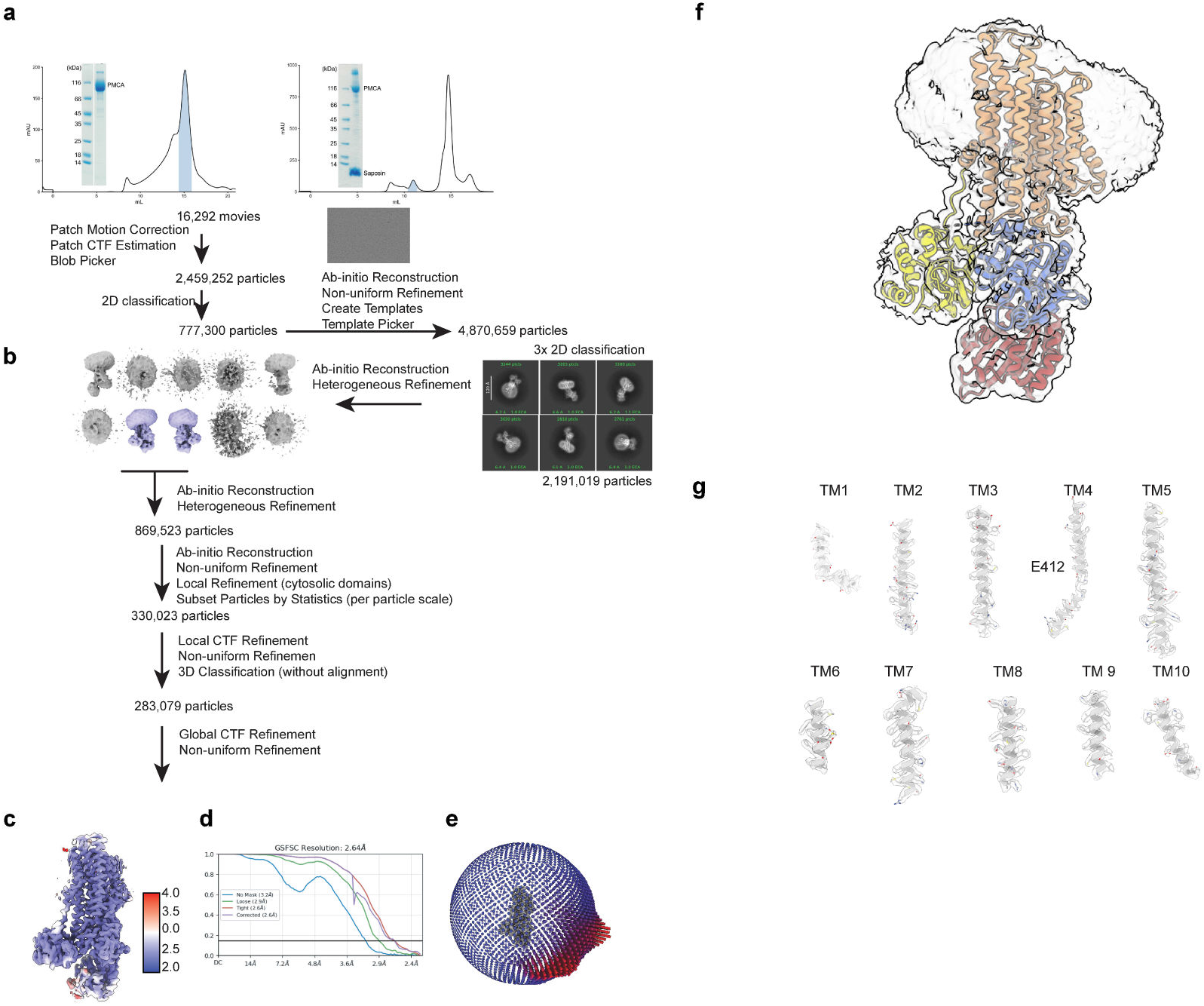
PMCA structure determination workflow. A, B. Chromatogram and corresponding PMCA monomer fraction on SDS-PAGE from size exclusion chromatography of PMCA in DDM (A) or Salipro nanodisc (B). C. Cryo-EM data processing workflow including representative micrograph, 2D classes, and Heterogeneous Refinement output volumes. D. PMCA map colored by local resolution. E. Fourier Shell Correlation (FSC) curves between half maps from final 3D reconstruction. F. Euler angle distribution on particles used in the final 3D reconstruction. G.Sample EM densities for the 10 transmembrane helices of human PMCA2z/a in an E1 Ca^2+^-bound conformation. All helices are depicted at a contour level of 0.15 (ChimeraX).

**S12.**
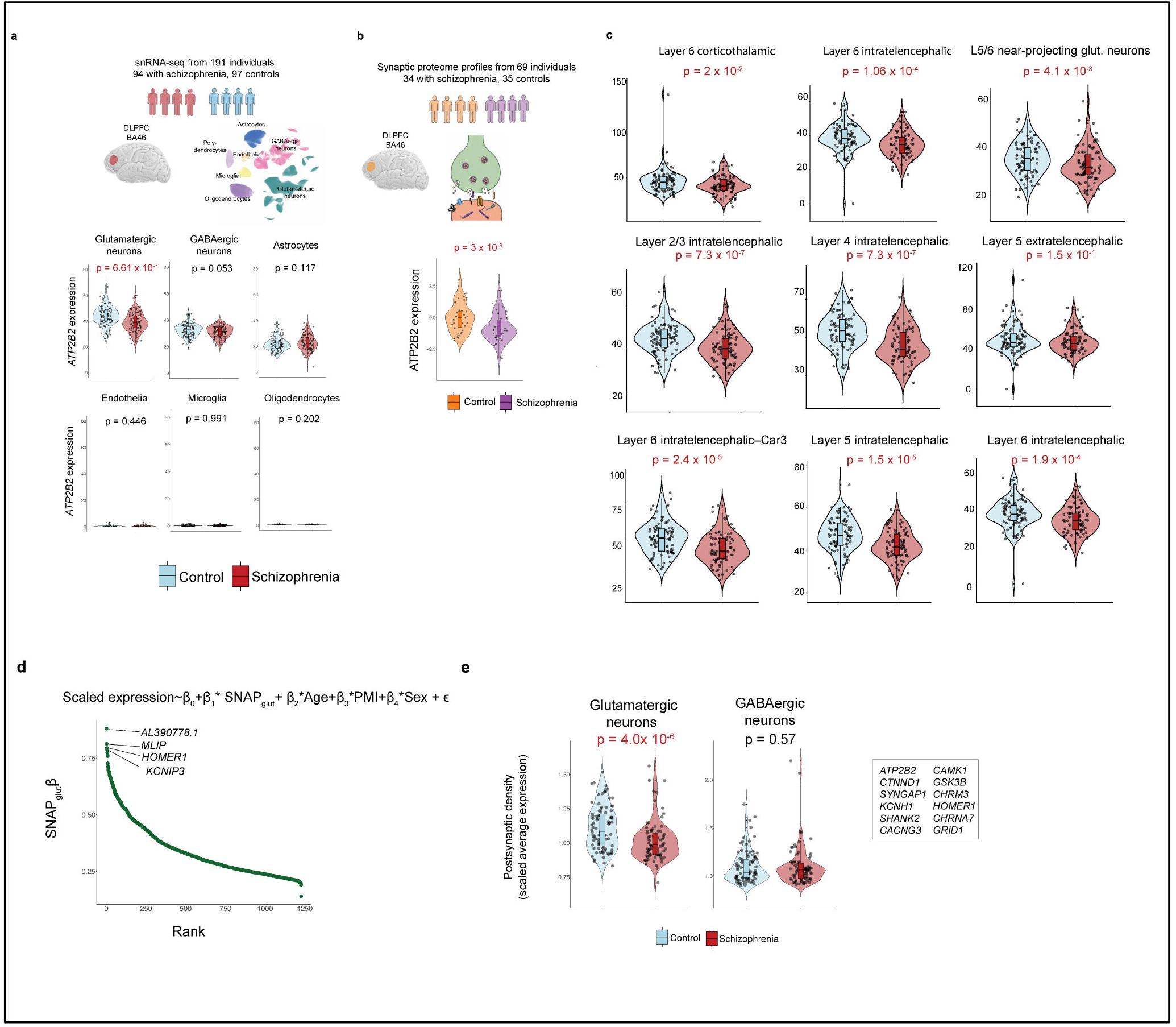
Expression of ATP2B2 in donors with and without schizophrenia. A. *ATP2B2* expression (per 10^5^ detected nuclear transcripts) comparison in persons with and without schizophrenia (n = 93 controls, 87 cases) across glutamatergic neuron subtypes of distinct cortical layers. Sequencing data obtained from 23 P-values are from a two-sided Wilcoxon rank-sum test. Box plots show interquartile ranges; whiskers, 1.5x the interquartile interval; central lines, medians; notches, confidence intervals around medians. B. ATP2B2 protein abundances are significantly under expressed in synapses in the DLPFC of donors with schizophrenia versus donors without. Shown is covariate-adjusted ATP2B2 synaptic protein abundance measurements generated in schizophrenia cases and controls. P-values are computed from a linear regression model (Methods). Box plots display interquartile ranges, whiskers extend to 1.5 times the interquartile range, central lines represent medians, and notches indicate confidence intervals around the medians. C. *ATP2B2* expression (per 10^5^ detected nuclear transcripts) comparison in persons with and without schizophrenia (n = 93 controls, 87 cases) across glutamatergic neuron subtypes of distinct cortical layers. Sequencing data obtained from^76^ P-values are from a two-sided Wilcoxon rank-sum test. Box plots show interquartile ranges; whiskers, 1.5x the interquartile interval; central lines, medians; notches, confidence intervals around medians. D. Expression of *ATP2B2* is positively correlated with a latent factor program, synaptic neuron and astrocyte program (Figure SNAP). Plotted is RNA expression of *ATP2B2* as a function of the donor loadings of SNAP in glutamatergic, GABAergic and astrocytes. Ranked coefficients of all significant genes from the regression of glutamatergic neuron expression on SNAP donor loadings. Genes specifically regulated by SNAP in glutamatergic, but not GABAergic, neurons are enriched for synaptic and postsynaptic density components, and exhibit coordinated downregulation in the glutamatergic transcriptomes of individuals with schizophrenia. P-value is from regression the average expression of core genes unto schizophrenia status (controlling for age, PMI and sex). Box plots show interquartile ranges; whiskers, 1.5x the interquartile interval; central lines, medians; notches, confidence intervals around medians. E. Average expression of postsynaptic density genes was computed per sample and compared between schizophrenia cases and controls in glutamatergic and GABAergic neurons. In glutamatergic neurons, cases show reduced expression relative to controls (P = 4.0 × 10^−10^)_2_ whereas no significant difference is observed in GABAergic neurons (P = 0.57). Points represent individual samples; violin plots show the distribution with embedded boxplots. Select genes are shown on the right.

### 3D neighborhood analysis

We mapped all 331 variants from SCHEMA and ASD onto the AlphaFold3-predicted structure of ATP2B2. To exclude poorly modeled regions, we restricted analysis to residues with high-confidence AlphaFold3 predictions (local pLDDT > 50 and PAE < 15 A), this resulted in the exclusion of 60 low-confidence residues and retaining 257 of 331 variants. For each unique residue harboring a variant, we defined a three-dimensional neighborhood by placing a sphere of radius 15 A around that position. Residue-residue distances were computed as the minimum interatomic distance between any atoms of the two residues, thereby capturing direct physical proximity including side-chain contacts, 106-109 (for comparisons of alternative radii, see Figure S7a). We then tested whether each 15 A spherical neighborhood was enriched for case relative to control variants using a Fisher’s exact test, comparing the case-to-control ratio within the neighborhood to that observed across the rest of the protein. Case variants exhibited higher predicted AlphaMissense pathogenicity scores than control variants (Figure S7f).

Missense variants were mapped to residues of an Alpha Fold predicted ATP2B2 structure. For residues *r* and *s*, the pairwise distance was defined as the minimum interatomic Euclidean distance:

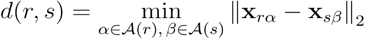

Residue pairs with low structural confidence were down-weighted by inflating distances when the predicted aligned error exceeded 15 A in both directions. Residues with mean pLDDT ~50 were excluded from analysis.

For each residue *r*, a three-dimensional neighborhood was defined as:

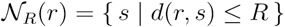

The number of case *c*_*r*_(*R*) and control *K*_*r*_(*R*) variants within the neighborhood of residue *r* were then given by

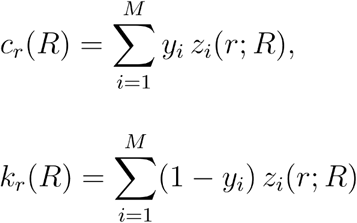

Let,

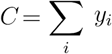 and 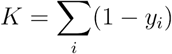 denote the global numbers of case and control variants, respectively. Enrichment of case variants in the spatial neighborhood of residue r was assessed using a one-sided Fisher’s exact test on the contingency table:

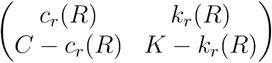

A Bonferroni correction adjusted the significance to account for the number of tested residues.

To identify independent clusters of missense variation, residues comprising Bonferroni-significant neighborhoods from the initial scan were removed, and the analysis was repeated on the remaining dataset using the same 3D neighborhood framework (15 A radius, Fisher’s exact test).

### JD neighborhood enrichment on PMCA2z/a cryo-EM structure

Missense variants defined on the canonical ATP2B2 sequence (Uniprot ID: **Q01814**) were remapped to the PMCA2z/a isoform and corresponding residue numbering derived from the cryo-EM model. Canonical and isoform sequences were globally aligned to generate a position mapping, and the isoform sequence and residue indices were extracted from the PMCA2z/a cryo-EM structure ^110^. A second alignment propagated isoform positions to structure-based residue numbers. Variants absent from the isoform or unresolved in the structure were excluded.

As with 3D neighborhood analysis on the AlphaFold3 model, residue-residue distances were computed from atomic coordinates in the PMCA2z/a cryo-EM model using minimum inter-atom distances between residues. Three-dimensional neighborhoods were defined as all residues within 15 A radius of each variant-containing position. Case and control variant counts within each neighborhood were compared to global counts using a one-sided Fisher’s exact test, with Bonferroni correction for multiple testing. Neighborhood-level p values were merged back onto the variant dataset.

### ScRNA-seq data pre-processing

For the caudate, we used data from ^111^ as previously reported, and DLPFC snRNA-seq data is from ^23^. We generated the amygdala dataset as per previously published protocols. Briefly, Frozen amygdala (N=4) and hippocampus (N=1) were obtained from the Harvard Brain Tissue Resource Center (HBTRC; McLean Hospital). History of psychiatric or neurological disorders was ruled out by consensus diagnosis carried out by retrospective review of medical records and extensive questionnaires concerning social and medical history provided by family members. All brain regions were examined by a neuropathologist. The cohort used for this study did not include any individuals with evidence of gross and/or macroscopic brain changes, or a clinical history of a cerebrovascular accident or other neurological disorders. Participants with Braak stages III or higher (modified Bielchowsky stain) were not included. None of the participants had a history of substance dependence within 10 or more years of death, as further corroborated by negative toxicology reports. Comprehensive health records, detailed family questionnaires, toxicology screening and neuropathology reports were reviewed to exclude donors with neurodegenerative or psychiatric disorders, as well as recent history of substance use. Donor metadata (age, sex, post-mortem interval) for these donors is included in **Supplementary Table 11**.

#### Single-nucleus library preparation and sequencing

The amygdala is from frozen tissue blocks from the amygdala of four healthy brain donors. For each donor, the full amygdala in its rostro-caudal, dorso-ventral, and medio-lateral extent was contained in two to three coronal blocks. These blocks were sectioned using a cryostat. Approximately every 100 µm, a section was mounted on a glass slide and stained with Luxol Blue to delineate the boundaries of the amygdala. These boundaries were then lightly etched (~ 100 µm depth) on the tissue block surface using a surgical blade to allow precise dissection of the amygdala in subsequent sections. Ten to twelve serial sections (12 µm each) were collected into Eppendorf tubes, ensuring that each tube contained a representative sampling from the rostral, middle, and posterior amygdala. One vial per case was used for RNA sequencing. For the temporal lobe, Nuclei were isolated from frozen tissue using a previously described protocol, which is publicly accessible at Protocols.io (https://doi.org/10.17504/protocols.io.4r3l22e3xl1y/v1). Briefly, frozen samples were processed to obtain clean nuclear suspensions suitable for downstream single nucleus applications. Single nucleus capture, gel bead in emulsion generation, and cDNA library construction were carried out using the 10x Genomics Chromium platform following the Chromium Next GEM Single Cell 3’ v3.1 chemistry guidelines (document version CG000204, Rev D), in accordance with the manufacturer’s instructions. After aligning to human reference GRCh38 with non-canonical contigs masked out, the libraries were run through CellBender to remove technical artifacts.

To address variability in sequencing depth across cells from all brain regions, we applied uniform normalization techniques to each count matrix. This involved normalizing the total number of unique molecular identifiers (UMIs) per cell, converting these counts to transcripts per 10,000 (TP10K), and then taking the logarithm to derive the final expression unit: log2(TP10K + 1). Seurat was employed for various analytical processes including data scaling, transformation, clustering and dimensionality reduction^112^. The SCTransform() function was used to scale and transform the data, as well as to identify variable genes^113^. Linear regression was performed to eliminate unwanted variations related to cellular complexity ( Such as the number of genes and UMIs per cell) or cell quality (percentage of mitochondrial and rRNA reads). Principal component analysis (PCA) was conducted using the identified variable genes, with the first 20 principal components (PCs) utilized to perform UMAP, embedding the dataset into two dimensions.

Subsequently, these first 20 PCs were used to construct a shared nearest-neighbor (SNN) graph via the FindNeighbors() function. This SNN was then leveraged to cluster the dataset using the FindClusters() function, implementing a graph-based modularity optimization algorithm based on the Louvain method for community detection.

### Gene set and drug target analyses

To broadly prioritize schizophrenia-relevant processes, we used GOrilla^47^, which identifies enriched GO terms in a given list of genes, to identify genes highly nominated for each phenotype by the prioritization analysis. We excluded terms that contained less than 10 genes. We used the Ensembl identifiers of the underlying genes from either analysis as foreground set and all genes used in the prioritization analysis as background. Analysis of gene sets specific to nervous system biology which have been previously implicated in neuropsychiatric disorders were also included, such as translational targets of FMRP^62^, chromatin targets of CHD8^63^, splice targets of RBFOX^64^ and *CELF4* target genes, (defined as genes with an iCLIP occupancy > 0.2)^114^.

For drug target analysis, we manually extracted gene sets from Open Targets^115^ for all eight neuropsychiatric disorders. For each disorder, we selected all genes with drug score > 0 (clinical trial phase 0 and above); each trait was well powered, containing at least 43 genes (anorexia) and up to 309 genes (epilepsy). For MAGMA, the per-gene p-values are calculated using the SNP2GENE function from FUMA for each trait, except for insomnia, which was extracted from Watanabe et al.^116^

### Ion Channel Gene Set Selection

To conduct ion channel analyses, we retrieved 215 ion channel genes from HGNC, grouped by major permeant ion (e.g; Ca^2+^, potassium, sodium, chloride). In addition to canonical channel types, we defined a composite category comprising volume-sensing channels, acid-sensing ion channels (ASICs), porins, and gap junction proteins (connexins and pannexins), reflecting their distinct functional roles and smaller representation in the genome; this group is labeled as “Volume & acid sensing, porin & gap junctions.” We restricted all analyses to genes that were expressed in the brain, defined by aggregating all frontal cortical neurons (excitatory and inhibitory) and selecting genes with logFC > 0, padj < 0.01, and avgExpr > 0.01 using the presto Wilcoxon AUC method^117^. Furthermore, within each ion channel class, we only tested genes that were also constrained (defined as having LOEUF observed/expected bin < 2). This filtered, neuronally expressed, and constrained gene set served as the background for all comparative analyses.

### Selective Constraint

We used LOEUF (loss-of-function observed/expected upper bound fraction, O/E) measures of intolerance to loss-of-function mutations extracted from gnomAD.v2’s pLoF metrics by gene data^15^. We used genes in the O/E bin of < 2 to denote constraint.

### Transcriptomic and proteomic schizophrenia case-control analysis

To assess the relationship between *ATP2B2* expression and schizophrenia status, we used snRNA-seq samples from ^23^, as provided by the authors, consisting of counts scaled to 100k UMIs per donor. We performed a multiple linear regression with ATP2B2 expression as the dependent variable, and schizophrenia status, age, sex, and post-mortem interval (PMI) as independent variables. We also confirmed that none of the schizophrenia, ASD, or NDD variants included in the 3D neighborhood analysis were present in this cohort.

For proteomic analyses, we used synaptic protein abundance measurements generated by mass spectrometry as in ^104^, which was log-transformed and processed through median centering as well as median absolute deviation scaling applied on a sample-by-sample basis. We constructed a linear model to examine the relationships between ATP2B2 protein abundance and covariates including diagnosis ( Schizophrenia versus control), age, sex, and post-mortem interval (PMI) across 69 individuals. To plot the effect of diagnosis controlling for confounding variables, we multiplied the matrix of covariates’ values (age, sex, and PMI) with their corresponding regression coefficients and subsequently adjusted the ATP2B2 levels by subtracting these offsets.

### GWAS summary statistics

We analyzed publicly available GWAS summary statistics for unique traits genetic correlation <0.9. We also used the summary statistics for SNPs with minor allele count >5 in a 1000 Genomes Project European reference panel ^118^.

### Burden Testing

For case-control analyses, we analyzed gene-level variant counts in autism from ^72^, bipolar disorder from ^119^, schizophrenia from^14^ major depression from^120^ and epilepsy from^121^ which included protein-truncating and missense variants. Genes were filtered for strong LoF constraint (LOEUF bin < 2) and merged with the prioritization-derived or MAGMA scores and the top 500 genes were selected. We summed the case and control counts of variants in these genes and tested for enrichment using a 2×2 Fisher’s exact test based on total variant counts and sample sizes. To assess whether our gene set exhibits stronger enrichment for rare damaging variants in autism cases compared to MAGMA, we performed a permutation-based burden test, where we compute the observed odds ratio for rare protein-truncating variants (PTVs) and missense variants with MPC > 3 in the top 500 prioritized genes using a 2×2 Fisher’s exact test. We then randomly sampled 500 genes from the same universe (all high pLI autism-relevant genes) 1,000 times and computed the OR for each random set. The empirical p-value was defined as the proportion of permuted gene sets with an OR greater than or equal to the observed OR.

For *de novo* burden analysis of rare variants in autism and neurodevelopmental disorders (Figure S**5**). We used the latest (unpublished) Autism Sequencing study (38,680 probands, 9,567 siblings) as well as *de novo* counts from neurodevelopmental disorders from ^38^. We assessed enrichment of *de novo* coding mutations by comparing observed mutation counts to expected rates based on gene-specific mutation models. For each gene, the observed number of *de novo* variants was derived by summing reported missense class 1 and class 2 mutations (defined by a combination of MPC and AlphaMissense scores), or PTVs, across all probands in the dataset. Expected mutation rates for each consequence class were obtained from published per-gene mutation rate estimates from gnomAD.

Expected *de novo* counts were computed by summing the mutation rates across the test gene set and scaling by twice the number of probands (to account for diploidy). Enrichment was calculated as the ratio of observed to expected mutations. Statistical significance was assessed using a two-sided Poisson test under the null hypothesis of no enrichment (rate ratio = 1). All analyses were performed separately for missense and PTV classes. Only constrained genes (LOEUF o/e < 2) were included in the analysis.

### In Vitro Measurement of ATPase Activity

#### Calmodulin production and purification

The pET15b-CaM (UniProt ID P0DP23) construct was transformed into chemically competent *E. coli* C41(F^−^, *ompT, hsdS*_*B*_(r_B_^−^ m_B_^−^), *gal, dcm* (DE3)) cells by heat shock. Positive transformants were selected on ampicillin-containing plates and grown in YP medium (10 g/L yeast extract, 16 g/L peptone, 5 g/L NaCl, 0.1 mg/ml ampicillin) until induction with 1 mM IPTG at an OD_600nm_ of 0.6. After 16 h at 21 °C, cells were harvested by centrifugation at 7,800 x g for 25 min at 4 °C, resuspended in a 1:2 w/v ratio in CaM resuspension buffer (20 mM Tris/HCl pH 7.5, 1 mM EDTA pH 8, and 5 mM β-mercaptoethanol) with protease inhibitors (1 mM PMSF, 1 µg/ml leupeptin, 1 µg/ml pepstatin A, and 1 µg/ml chymostatin), and disrupted by sonication. Cell debris was removed by centrifugation at 14,000 x g for 25 min at 4 °C, and the supernatant was mixed 1:1 with CaM wash buffer (20 mM Tris/HCl pH 7.5 and 10 mM CaCl_2_) and loaded onto a phenyl sepharose (Cytiva) column pre-equilibrated in CaM Wash Buffer. The column was washed sequentially with 5 cv each of (i) CaM Wash Buffer, (ii) CaM salt-wash buffer (20 mM Tris/HCl pH 7.5, 500 mM NaCl, and 10 mM CaCl_2_), (iii) CaM wash buffer. CaM was eluted in CaM elution buffer (20 mM Tris/HCl pH 7.5, 150 mM NaCl, and 10 mM EGTA) and subjected to size exclusion chromatography on a HiLoad 16/600 Superdex 75 pg column (Cytiva) in CaM SEC buffer (20 mM Tris/HCl pH 7.5 and 30 mM NaCl).

Purified CaM was used in ATPase activity measurements or for CaM-sepharose preparation. To prepare CaM-sepharose, purified CaM was subjected to buffer exchange on a PD-10 Desalting Column (Cytiva) to Coupling Buffer (50 mM HEPES/NaOH pH 8.3, 500 mM NaCl). For 120 mg of CaM, 5 g of lyophilized CNBr-Activated Sepharose 4B (Cytiva) resin was dissolved and washed in 1 L cold 1 mM HCl on a 0.45 µm cellulose acetate membrane filter (Labsolute). The resin was dried under vacuum, mixed with CaM, and incubated at rt for 1 h, before being loaded onto a gravity column and washed with 5 cv Coupling Buffer and 2 cv Blocking Buffer (100 mM Tris/HCl pH 8.0). The resin was mixed in Blocking Buffer at rt for 2 h and washed through three cycles of (i) 5 cv 100 mM acetate pH 4, 500 mM NaCl, and (ii) 5 cv 100 mM Tris/HCl pH 8, 500 mM NaCl. The resin was then washed in 5 cv water and stored in 20% EtOH at 4 °C until use in PMCA purification.

#### PMCA constructs

Codon-optimized PMCA2z/a (GenScript, UniProt ID Q01814-4) was cloned into a pEMBL-yex4 expression plasmid by homologous recombination in *S. cerevisiae* ^*122*^. The construct was subjected to site-directed mutagenesis using QuikChange Lightning Site-Directed Mutagenesis Kit (Agilent), Q5 Site-Directed Mutagenesis Kit (New England Biolabs), or Phusion Hot Start II High-Fidelity PCR Master Mix (Thermo Fisher Scientific). Primer design and site-directed mutagenesis workflow followed the manufacturers’ protocols. All constructs were verified by sequencing (Eurofins). See Supplementary Methods section **Mutagenesis primers for ATP2B2** constructs for the primers used.

#### Yeast transformation and PMCA production

PMCA expression vectors were introduced into calcium pump-depleted *S. cerevisiae* K616 (MATα *pmr1::HIS3 pmc1::TRP1 cnb1::LEU2, ade2, ura3, can1*)^123^ using the LiAc/ssDNA/PEG procedure^124^. Positive transformants were selected on uracil-deficient plates, grown in minimal medium (2% glucose, 40 µg/ml adenine hemisulfate, 10 mM CaCl_2_, 0.17% yeast nitrogen base w/o amino acids and w/o ammonium sulfate, 0.5% ammonium sulfate, and 0.08% CSM-Ura w/Ade40 (Sunrise Science Products)) at 30 °C, and induced with 2% galactose, 1% yeast extract, and 2% peptone at an OD_600nm_ of ~5. After 18 h at 18 °C, cells were harvested by centrifugation at 1000 x g for 15 min at 4 °C, washed by resuspension and centrifugation at 1000 x g for 15 min at 4 °C in (i) 1:3 w/v water and (ii) 1:2 w/v TEKS buffer (50 mM Tris/HCl pH 7.6, 2 mM EDTA, 100 mM KCl, and 0.6 mM D-sorbitol). The resulting pellet was resuspended 1:1 w/v in TES buffer (50 mM Tris/HCl pH 7.6, 2 mM EDTA, 0.6 mM D-sorbitol, and 5 mM β-mercaptoethanol) with protease inhibitors.

#### PMCA purification

All purification steps were performed at 4 °C. Cells were disrupted with 0.5 mm glass beads in a BeadBeater (BioSpec Products) through 6 cycles of 1 min disruption / 2 min break. Unbroken cells were removed by centrifugation at 2,000 x g for 20 min, and the supernatant was centrifuged at 20,000 x g for 25 min to give S2, which was then ultracentrifuged at 136,000 x g for 3 h. The resulting membrane pellet was resuspended 1:5 in resuspension buffer (50 mM Tris/HCl pH 7.2, 150 mM KCl, 20% glycerol, and 5 mM β-mercaptoethanol) with protease inhibitors, homogenized in a Dounce glass homogenizer, flash-frozen in liquid N_2_, and stored at −80 °C.

Membranes were solubilized in 1% n-dodecyl-β-D-maltoside (DDM) and 0.2% cholesteryl hemisuccinate (CHS) for 1 h, followed by ultracentrifugation at 177,000 x g for 30 min. The supernatant was supplemented with 4 mM CaCl_2_ and incubated with CaM Sepharose beads for 1 h. Beads were washed sequentially with 10 cv each of (i) binding buffer (50 mM Tris/HCl pH 7.2, 150 mM KCl, 2 mM CaCl_2_, 10% glycerol, and 0.017% DDM), (ii) high-salt buffer (Same composition, 250 mM KCl), and (iii) binding buffer. Protein was eluted with elution buffer (50 mM Tris/HCl pH 7.2, 250 mM KCl, 2 mM EGTA, 10% glycerol, and 0.017% DDM), concentrated to 3-5 mg/ml in a 100 kDa MWCO Amicon® Ultra-4 Centrifugal Filter unit (Merck Millipore), and subjected to size exclusion chromatography on a Superose 6 increase 10/300 column (GE Healthcare) in SEC buffer (50 mM Tris/HCl pH 7.2, 150 mM KCl, 10% glycerol, and 0.017% DDM). Monomeric fractions were pooled, re-concentrated to 1-2 mg/ml, flash-frozen in liquid N_2_, and stored at −80 °C until use.

#### ATPase activity assay

Phosphatidylcholine (PC) stocks for relipidation were prepared as follows: Powdered egg PC (Avanti Research) was dissolved in chloroform, dried to a thin film under N_2_ gas, and resuspended to 10 mg/ml in 50 mM Tris/HCl pH 7.2, 150 mM KCl, 1.5% DDM. A translucent gel was obtained after cycles of vortexing, sonication, and heating to 50 °C. Prior to ATPase activity measurements, purified PMCA was relipidated in PC in a 1:5 w/w ratio for 15 min, after which the protein-lipid mixture was added to a final protein concentration of 10 µg/ml to a reaction mixture (40 mM BisTris/HEPES pH 7.2, 3 mM MgCl_2_, and 1 mM EGTA) ^125^. Ca^2+^, CaM, and/or corresponding buffers were added to desired concentrations, and the concentration of free Ca^2+^ was calculated using the Maxchelator Ca/Mg/ATP/EGTA Calculator v1.0 with constants from (Schoenmakers et al., 1992). Reactions were initiated by addition of 3 mM ATP and stopped after 14 min at 37 °C by mixing 50 µl reactions with 50 µl colorimetric solution (28.3 mM ammonium molybdate VI tetrahydrate mixed 1:5 with 0.17 M ascorbic acid, 0.1% SDS in 0.5 M HCl). After 10 min, 75 µl arsenic solution (20 mg/ml NaAsO_2_, 20 mg/ml trisodium citrate 2xH_2_O, and 2% acetic acid) was added to stabilize the solution and prevent reaction between molybdate and newly released phosphate^126^. Absorbance was measured at 860 nm and compared to a phosphate standard.

#### Functional complementation assay for PMCA Ca^2+^ transport

An empty pEMBL-yex4 expression vector or pEMBL-yex4 expression vectors containing PMCA variants under control of a galactose inducible promoter were transformed into K616 cells as previously described. Positive transformants were taken directly from the uracil-deficient selection plates, suspended in sterile H_2_O, and diluted to a starting OD 600nm of 2, 1,0.5, 0.2, 0.1, 0.01, and 0.001. 5 µl of each dilution step was dropped onto plates containing 2 % galactose or 2 % glucose with either 10 mM CaCl_2_ or 10 mM EGTA. The plates were incubated for 72 h at 30 °C before imaging. As the K616 yeast strain lacks the two endogenous Ca^2+^-ATPases PMR1 and PMC1, it is unable to grow on Ca^2+^-depleted media. However, expression of an active Ca^2+^-pump can complement the strain, presumably by scavenging Ca 2+ from the cytosol to the secretory pathway ^127^. Only cells expressing an active PMCA variant were therefore able to grow on Ca^2+^-depleted media in the complementation assay.

#### Cryo-EM sample preparation and data collection

PMCA was reconstituted in Salipro® nanoparticles before cryo-EM data collection ^128^. PC stocks were diluted to 5 mg/ml in Salipro buffer (50 mM Tris/HCl pH 7.2 and 150 mM KCl) and mixed with PMCA and saposin A in a 1/1.7/4.4 (PMCA/lipids/saposin) w/w/w ratio. The mixture was diluted in Salipro buffer to reduce the DDM concentration below the critical micelle concentration and was thereafter concentrated in a 100 kDa MWCO Amicon® Ultra-4 Centrifugal Filter unit (Merck Millipore) prior to size exclusion chromatography on a Superdex 200 increase 10/300 column (Cytiva). Peak monomer fractions were pooled and concentrated to 0.5 mg/ml. 3 µl sample was applied to glow discharged (60 s, 15 mA) Quantifoil® R 1.2/1.3 300 Mesh Cu grids. The grids were blotted for 5.5 s at blot force 0 and plunged into liquid ethane in a Vitrobot Mark IV (Thermo Fisher Scientific) operated at 4 °C and 100% humidity. Grids were stored in liquid nitrogen until data collection.

Movies were collected on a Titan Krios G3i transmission electron microscope (Thermo Fisher Scientific) equipped with a K3 direct electron detector (Gatan) at 130,000x magnification, giving a pixel size of 0.647 Å, using EPU 3.8.1 software (Thermo Fisher Scientific). A nominal defocus value range of 0.6-2.0 µm was used at a total dose of 59.9 e^−^/Å^2^ in 53 frames with an exposure time of 1.4 s per movie. 16,292 movies were collected in total. The data collection parameters are summarized in **Supplementary Table 11**.

#### Cryo-EM data processing and model building

The cryo-EM dataset was processed in CryoSPARC v4.7.0 ^129^. The collected movies were corrected for beam induced motion by Patch Motion Correction, and the CTF (contrast transfer function) was estimated by Patch CTF. Micrographs unsuitable for data processing were removed by manual inspection. Particles were initially picked by a Blob Picker with a size range of 100 to 180 Å, and extracted particles were sorted by 2D Classification followed by Ab-initio Reconstruction of a 3D volume. The best class was selected for Non-uniform Refinement, and the resulting volume was used to generate 2D templates for Template Picking.

The newly picked particles were sorted by three rounds of 2D Classification followed by two rounds of Ab-initio Reconstruction and Heterogeneous Refinement. A consensus volume was generated with Non-uniform Refinement, and the density of the cytoplasmic domains was improved by Local Refinement using a mask around the N, P, and A domains. The particles were then sorted based on their per-particle scale using two Gaussians. The resulting particle stack was used for Local CTF Refinement followed by final sorting by 3D Classification without alignment. Two rounds of Global CTF Refinement were used to fit beam tilt, anisotropic magnification, and spherical aberrations. The final 3D volume was then generated by Non-uniform Refinement, and the local resolution was estimated. An overview of the data processing workflow is presented in figure x.

The initial atomic model was generated with AlphaFold 3 ^37^. Using ChimeraX 1.10 ^130^ the unresolved regions were trimmed, and each domain was disconnected from the model and fitted to the density map separately. The model was then put back together and iteratively refined using ISOLDE ^131^, Phenix ^132^, and Coot ^133^. Figures were prepared using ChimeraX 1.11 ^130^ and PyMol.

#### Functional testing of ATP2B2 variants in HEK cells using GCaMP6s-CAAX

##### Cell culture

HEK293T/17 cells were used for generation of lentivirus, and grown in DMEM supplemented with 10% FBS, 100 U/ml penicillin and 100 µg/ml streptomycin. HEK293 CaV3/Kir2.3/GCaMP6 cells were grown in DMEM/F12 media supplemented with 10% FBS, 100 U/ml penicillin and 100 µg/ml streptomycin, and were cultured as previously described^91^. All cells were grown at 37 °C, 95% humidity with 5% CO2.

##### Lentivirus production

Lentivirus was produced by co-transfecting HEK293T/17 cells with the desired transfer plasmids (or pooled library) and two packaging vectors (psPAX2 and pMD2.G, Addgene #12260 and #12259) using Fugene HD transfection reagent. 6-12 hours post transfection, fresh media was added to the HEK cells, and virus was collected through collecting the media supernatant at 48 hours post transfection then flash frozen. Virus was rapidly thawed prior to transduction.

##### Plasmids and cell line generation

gblocks containing the WT or E457K ATP2B2 sequences were purchased from Integrated DNA Technologies (IDT) and cloned into a lentiviral expression vector under the PGK promoter, generating constructs with an mCherry-P2A sequence upstream of either the WT or E457K ATP2B2 coding sequence. Lentivirus was produced from each construct and used to transduce HEK293 CaV3/Kir2.3/GCaMP6 cells at a multiplicity of infection (MOI) of 0.1 by adding the virus to the media following cell splitting. Given the potential for lentiviral transgene silencing in HEK cells, transduced cells were sorted based on mCherry expression (as a proxy for ATP2B2 expression) one day before seeding for the FLIPR assay.

##### FLIPR assay for testing calcium levels

A day after sorting cells based on mCherry expression, 15,000 cells per well were seeded into poly-D-lysine-coated 384-well clear-bottom plates with 1 µg/mL doxycycline. Each plate included 128 wells per condition: (1) mock WT HEK293 CaV3/Kir2.3/GCaMP6 cells, (2) HEK293 CaV3/Kir2.3/GCaMP6 cells expressing WT ATP2B2, and (3) HEK293 CaV3/Kir2.3/GCaMP6 cells expressing E457K ATP2B2. Two test plates were used, along with a KCl titration plate to determine the EC30 and EC90 values for the assay. Two days after plating, the FLIPR assay was performed as previously described^91^.

## Calcium Imaging and Quantification of ATP2B2 Variant Function

To enable comparison of ATP2B2 variant effects across experiments, we applied a model based analysis of calcium response magnitude to isolate variant specific effects while controlling for plate to plate differences. Cells expressing A TP2B2 variants or wild type controls were plated into *t*_1_ = 40 FLIPR plates and a time series fluorescence was collected at 1 Hz across all wells and plates. A single depolarizing stimulus was delivered using a KCI addition at time (in seconds), and raw data consisted of fluorescence traces *F*_*i,p,w*_(*t*) for variant *i*, plate *p*, well *w*, and time t. We performed well specific baseline normalization. For each plate and well, the baseline fluorescence was defined as the mean signal prior to the KCI stimulus. Baseline corrected fluorescence was computed as;

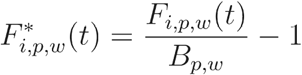

where *B*_*p,w*_ is the mean fluorescence in each well before the KCI stimulus. The calcium response following KCI stimulation was quantified as the area under the baseline corrected fluorescence curve. For each well, the AUC was calculated via integration using the using the trapezoidal rule,

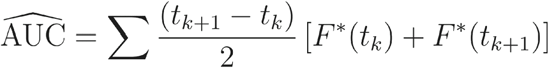

To estimate variant specific differences while accounting for plate variability, we fit the linear mixed effects model:

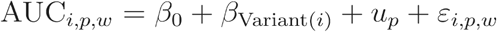

with wild-type as the reference level. The random 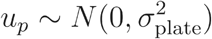 intercept accounts for plate specific shifts in *ε*_*i,p,w*_ overall AUC, and captures residual variation. Parameters were fit using restricted maximum likelihood in the lme4 R package ^134^. Estimated marginal means were then calculated to summarize the model adjusted mean AUC for each variant after accounting for plate effects ^135^. Differences between each variant and wild-type were then computed, and the confidence intervals are calculated from the estimated difference between each variant and wild-type and the standard error of that estimate, which incorporates both within plate variability and plate to plate variability from the mixed model.

### Western Blot Sample Prep and Detection

Cells were collected and lysed by rocking them for 15 minutes at 4 °C in RI PA lysis buffer (Thermo) supplemented with 1x Halt Protease Inhibitor Cocktail (Thermo Fisher, 78429), followed by centrifugation for 5 minutes at 5,000 ref to obtain clarified cell lysates. Protein concentrations within the lysates were quantified using the Pierce BCA Protein Assay kit (Thermo Fisher, 23225) for equal loading. 6x Laemmli SOS buffer was then added to 6 µg of protein that were then either run directly, or first boiled at 95°C for 5 minutes before proteins were separated on Bolt 4-12% Bis-Tris gels (Thermo Fisher, NW04127BOX). After the run, proteins on the gel were transferred to Nitrocellulose membranes using the Mini Trans-Blot Cell (BioRad) kit at the high molecular weight settings. Following transfer, membranes were blocked with EveryBlot Blocking Buffer (BioRad, 12010020) at room temperature for 1 hour, and subsequently incubated with primary antibodies, at dilutions indicated by the manufacturer, overnight at 4°C. The following day, membranes were washed 3 times with 1X TBST followed by a 1 hour incubation with the secondary antibodies. 3 washes in 1X TBST were then performed and the membranes were then incubated with 2 ml working solution of SuperSignal West Pico PLUS Chemiluminescent Substrate (Thermo, 34577) and imaged using a ChemiDoc MP system (Biorad). The following antibodies were used: Primary antibodies against ATP2B2 (abeam, ab3529), alpha-tubulin (abeam, ab7291), mCherry (Proteintech, 26765-1-AP) and GAPDH (Cell signaling technology, 97166). Secondary antibodies used were anti-rabbit lgG, HRPlinked (CST 7074) and anti-mouse lgG, HRP-linked (CST 7076).

### Simulating Variant Effects on Per-Residue Contact Probabilities

AlphaFold3 can model protein-ligand interactions, such as ATP2B2 with Ca^2+^. Ca^2+^ is represented as a single-atom ligand token under the AF3 tokenization scheme (where protein sequences are tokenized as residues, nucleic acids as nucleotides, and each ligand or ion becomes its own token). Its atomic features (element identity, formal charge, reference conformer coordinates, atom-name embedding, and reference-space identifiers) are incorporated into the atom-level representation through the AtomAttentionEncoder, which aggregates this atomic metadata to produce a single vector that serves as the model’s internal representation of the Ca^2+^. Token-level embeddings are generated for both protein residues and the Ca^2+^ ion. For residues, the embedding integrates atom-derived features with residue-type information and MSA-derived conservation and deletion statistics. For Ca^2+^, which is represented as a single-atom ligand token, the embedding is derived solely from its *z*_*init*_[*i*, Ca] atomic features.

For each residue-calcium pair (*i, Ca*), the model constructs an initial pair embedding using (i) a linear projection of the single-token embeddings of residue i and the calcium ion, and (ii) a learned relative positional encoding that incorporates token index offsets, chain identity, and residue indexing. This representation encodes chemical compatibility, coarse spatial priors, and topological structure implied by the input features.

The pair embedding is then refined by the Pairformer stack, which applies 48 sequential blocks of triangular multiplicative updates and triangular self-attention over all token triplets (*i, k*, Ca,). These updates propagate geometric constraints such that inferred proximities can reinforce one another: for example, if residue -i is predicted to be close to residue *k*, and residue *k* is *z*_*init*_[*i*, Ca] predicted to interact with Ca^2+^, then the model adjusts accordingly. After recycling through these updates, the refined pair embeddings *z*_*trunk*_[*i*, Ca] represent AF3’s latent, geometry-aware estimate of the residue-ion spatial relationship, independent of the final coordinate sample produced by the diffusion module.

These refined pair embeddings are passed to the distogram head, which projects each *z*_*init*_[*i*, Ca] onto 64 distance bins spanning approximately 2.3-21.7 A. A softmax applied to the resulting logits yields a categorical probability distribution

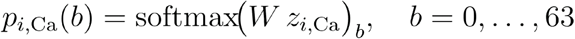

where each bin corresponds to a fixed interval [*d*_min_ (*b*), *d*_max_(*b*)). A residue–calcium contact probability at threshold *d*_*c*_ is defined as the summed probability mass over all bins whose upper boundary lies below the cutoff:

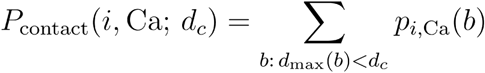

where a conventional definition of contact is 8 Å ^136^. For each ATP2B2 model (wild type and variants), we parsed the AlphaFold 3 JSON output from alphafoldserver.com and extracted the contact_probs matrix, which contains the predicted probability that any residue-token pair lies within 8 A. Because Ca^2+^ is represented as a single-atom ligand token, its corresponding column in this matrix (column 1244 in our runs) reports the model’s estimated proximity between each residue and the ion. The same procedure was applied to the variants near the ATP:Mg^2+^-binding regions.

### Electrostatic potential calculations

All structural analyses began with an AlphaFold3 model of human A TP2B2, which served as the common template for all missense variant structures. Individual missense variants were introduced using FOLDX v5 ^87^. The workflow used the standard RepairPDB followed by BuildModel sequence. RepairPDB was applied once to the AlphaFold template, either the model used in complex with Ca^2+^ or ATP:Mg^2+^, BuildModel was run with default settings for each mutation. No custom force field options, backbone constraints, or nonstandard parameters were used.

Each FOLDX generated structure was converted to a PQR file using PDB2PQR v3.7.1 ^137,138^. The conversion was performed with the PARSE force field and PROPKA based titration states at pH 7. Electrostatic potentials were computed with Adaptive Poisson-Boltzmann Solver (APBS). All runs used identical grid dimensions, grid spacing, dielectric constants, and solvent settings. This produced OpenDX (.dx) potential maps for wild type and all mutant structures under matched numerical conditions.

To quantify substitution induced changes in the local electrostatic environment, we created a Python workflow that loads both the PQR coordinates and the APBS potential grids, and uses the geometric center of the mutated residue in the wild type structure and uses this point as the center of a 5 A spherical region on the APBS grid. The wild type and mutant potential maps were parsed from the APBS .dx files, and grid consistency was confirmed before analysis. For each variant, all grid points within a 5 A sphere centered on the mutated residue (defined in the wild type structure) were extracted. The mean electrostatic potential in this region was computed for wild type and mutant, and the difference was used as the mutation’s local perturbation score. Variants were annotated as case or control and ranked by the magnitude of their local potential shift.

### LDSC

S-LDSC assesses the contribution of a genomic annotation to the heritability of a trait. Namely, it assumes that the per-SNP heritability or variance of effect size (of standardized genotype on trait) of each SNP is equal to the linear contribution of each annotation:

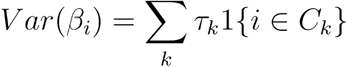

Where the marginal Chi-square association statistic for SNP *i* reflects the causal contributions of all SNPs in LD with SNP *i*.

Where a is a constant that reflects sources of confounding and N is the GWAS sample size ^40^, and ℓ(*i, k*) is the LD score of SNP; to category Ck. The regression coefficient τ_*k*_ quantifies the importance of annotation *C*_*k*_, correcting for all other annotations in the model. Specifically, *τ*_*k*_ which is the proportionate change in per-SNP heritability associated with a standard deviation change in the value of the annotation, conditional on other annotations included in the model ^41,42^.

### Transcriptomic analysis of neurons

Given a dataset of neuronal cell types, we first computed an expression estimate for each gene within a cluster (42 clusters in total) to reduce the influence of sampling variability and cell-to-cell heterogeneity. For gene *g*, let the mean expression across all cells be *µ*_*g*_, the standard deviation be *σ*_*g*_, and the number of cells in the cluster be *n*_*c*_. The standard error of the mean was defined as:

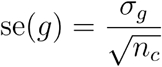

We then defined the adjusted expression value as:

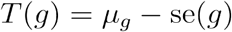

For each pair of cell types, we defined two one-sided differential expression gene sets: the top 1,000 genes enriched in cell type A relative to cell type B, and conversely the top 1,000 genes enriched in B relative to A. Genes were ranked by the 1092 fold-change of *T*(*g*), and we applied a floor on expression to exclude genes supported by very few reads.

All of these pairwise-defined gene sets were then used as annotations in stratified LO score regression to compute *τ*_*k*_, (conditioning on the full baseline model). We applied family-wise error rate correction across all tested pairs. We then defined an indicator function:

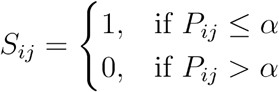

where *P*_*ij*_ is the raw p-value and *α* is the FWER-adjusted threshold.

For each gene, we consider all gene sets *G* in which *g* appears. From stratified LD score regression, each gene set *G* has an estimated per-SNP heritability enrichment coefficient *τ*_*G*_ and an associated standard error *se*(*τ*_*G*_). we then assign each gene a cumulative score equal to the sum of LOSC z-statistics across heritability-enriched neuronal contrasts in which that gene participates, restricted to those gene sets that survive FWER correction.

## Graphics

We used R to generate all plots (R version 4.1, 4.2 and 4.3). We generated enrichment heatmaps, gene term enrichment, error plots, box plots, distribution plots and scatterplots using a combination of ggplot2 (v.3.3.6) and ggpubr (v.0.4.15).

## Statistics and reproducibility

All data used in the present study were generated and designed by the original studies and no statistical method was used to predetermine sample size. No data were excluded from the analyses. The experiments were not randomized. The Investigators were not blinded to allocation during experiments and outcome assessment.

## Data availability

The cryo-EM density map has been deposited in the Electron Microscopy Data Bank under accession code EMD-56546. The corresponding atomic coordinates have been deposited in the Protein Data Bank under accession code 28JP. Raw cryo-EM movies will be deposited in EMPIAR and made publicly available upon publication. Single-nucleus RNA sequencing data from human amygdala and temporal lobe are available at Zenodo (https://doi.org/10.5281/zenodo.20024150). Amygdala data are provided as raw gene-by-cell count matrices, and temporal lobe data are provided as cell-type-level pseudobulk count matrices. Raw read-level and cell-level data for the temporal lobe sample are not deposited because participant consent for this archival dataset does not permit release of data from which donor genotype information could be recovered.

All other data supporting the findings of this study are available from the corresponding author upon reasonable request.

## Code availability

The 3D neighborhood test is available as an open-source Python package at *https://github.com/sherifgerges/3DNT_ATP2B2*.

## Supplementary Tables

<u>Supplementary Table 1. Numerical results of burden testing.</u>

Numerical results from a rare variant burden on prioritized genes (pertaining to Figure 1d).

<u>Supplementary Table 2. Numerical results of drug target testing.</u>

Numerical results from a Fisher’s test on drug target genes sourced from Open Targets (pertaining to S3b).

<u>Supplementary Table 3. Gene ontology analysis.</u>

Gene set enrichment results from prioritization analysis based on the top 500 ranked genes per phenotype.

<u>Supplementary Table 4. List of Ca2+ flux pathway genes.</u>

Full Ca^2+^ flux pathway gene list (related to Figure 2d) with corresponding P-values from exome sequencing studies for schizophrenia, ASD, NDD, DD, and Epilepsy.

<u>Supplementary Table 5. ATP2B2 variants associated with schizophrenia, ASD and NDD.</u>

Rare missense variants identified in case and control cohorts and de novo studies of ASD and NDD. For each variant, the predicted functional consequence, allele counts in cases and controls, AlphaMissense pathogenicity score, and associated phenotype are provided. Where applicable, structural annotations corresponding to calcium or ATP binding proximity are included. In the schizophrenia case-control analysis, only variants present either exclusively in cases or controls were used.

<u>Supplementary Table 6.</u>

Association statistics for the Ca^2+^ flux pathway gene list across neurodevelopmental and psychiatric disorders. For each gene, the Ensembl gene ID, LOEUF constraint bin, and P values from analyses of schizophrenia, neurodevelopmental disorder, autism spectrum disorder, and epilepsy are shown. Epilepsy results reflect protein truncating variant enrichment

<u>Supplementary Table 7.</u>

Ranked prioritization gene list with TauSum scores.

<u>Supplementary Table 8.</u>

Baseline-normalized calcium or fluorescence signal over time, summarized by mean and standard error per condition.

<u>Supplementary Table 9.</u>

Each row lists a missense variant, its amino acid position, and the corresponding per residue P value from the 3D neighborhood test (related to Figure 3b). These values are derived from the initial, unconditional analysis and reflect spatial enrichment of case variants within a 15 Å radius around each residue. P values were calculated using a one-sided Fisher’s exact test comparing the case to control ratio within the local neighborhood to that across the full protein.

<u>Supplementary Table 10</u>

Summary of microscope settings, image acquisition parameters, particle numbers, map resolution, refinement procedures, model composition, and structural validation metrics for the

2.64 Å cryo-EM structure of ATP2B2. Validation statistics include geometric quality measures, MolProbity and clash scores, Ramachandran and CaBLAM analyses, and deposition identifiers for the atomic model and EM density map.

<u>**Supplementary Table 11.**</u>

Metadata for human brain samples

## Supplementary Discussion

### Extended discussion of Ca^2+^ flux pathway gene selection

In defining our core set of “Ca^2+^ flux pathway genes” we intentionally focused on genes encoding calcium-selective channels excluding other elements of the neuronal Ca^2+^-signaling toolkit. Notably, we omitted N-methyl-D-aspartate-type receptors (NMDARs) and AMPA- and kainate-type glutamate receptors. While these receptors play a crucial role in transmitting calcium signals to the nucleus, triggering action potentials, and leading to plasma membrane depolarization and have roles in excitation-transcription coupling (reviewed in ^101^),, these receptors are non-selective channels..

### Extended Discussion: Simulating Mutation Effects on Per-Residue Contact Probabilities with AlphaFold3

These predictions provide a quantitative basis for identifying calcium-coordinating residues, and allows us to simulate the effects of substituting individual residues to gauge the change in per-residue contact probabilities. To further explore the structural and functional consequences of the E457K mutation, we revisited the underlying biophysical principles that might explain its disruptive effects. As previously noted, E457K (Figure 3f) induces a marked reduction in per-residue contact probabilities, which could be attributed to the substitution of a negatively charged glutamic acid with a positively charged lysine. To systematically investigate this hypothesis, we systematically mutated a glutamic acid (E) residue in the ATP2B2 protein to every other amino acid and measured the per-residue change in calcium contact probabilities (compared to wild type). This was quantified as the difference (residual) in contact probability. A larger residual reflected a larger perturbation between calcium ions and contact sites in ATP2B2.

As expected, mutations to Proline, Arginine, and bulky hydrophobic residues (e.g; Isoleucine, Valine, Methionine) caused the largest disruptions in calcium contact probabilities (Figure S6). These likely disturb local protein structure or calcium-binding sites due to charge reversal or steric clash. Reassuringly, mutations to chemically similar or small residues (e.g., Aspartic acid, Glutamine, Glycine) showed minimal disruption, suggesting partial retention of the wild-type interaction. These conservative substitutions produced minimal changes in predicted structure and calcium-binding behavior, suggesting that AF3 reasonably captures the structural impact of substitutions based on core biochemical properties. Taken together, these results suggest that substitutions that alter charge, polarity, or hydrophobicity tend to cause the most pronounced structural disruptions, whereas substitutions that retain the fundamental properties of glutamic acid result in comparatively minor perturbations.

### Extended discussion of limitations

This analysis uses stratified LD score regression (S-LDSC) to leverage cell-type–specific expression and the power of GWAS to detect genetic variation that modulates cellular processes in the brain. Its first limitation is the scope of tissues and the diversity of cell types and states represented. Second, the approach relies on identifying highly expressed gene programs, which may not reflect biological causality; if a causal cell type is not assayed, co-expressed genes and cell types may be implicated instead. However, such signals may still be informative for drug development by highlighting relevant pathways. Third, the analysis does not incorporate cell-type–specific eQTL data, which may better capture underlying regulatory mechanisms. Fourth, it does not distinguish whether implicated gene processes represent conditionally independent signals. Fifth, the LD score regression framework is primarily applicable to common and low-frequency variants, and less applicable to rare variant enrichments (we instead use rare variants to validate our findings). Sixth, we have focused on human scRNA-seq data; however, incorporating data from other modalities such as ATAC-seq (Assay for Transposase-Accessible Chromatin using sequencing, ^139^) and DNase-seq (DNase I hypersensitive sites sequencing ^140^) can more specifically map candidate cis-regulatory sequences. In turn, relevant animal models could allow experimental validation of neuropsychiatric disorders mechanisms in model organisms ^141^. Seventh, we identify programs by gene co-variation using only healthy tissues. Future work could utilize disorder relevant tissue to nominate more precise sets of genes and the cell types in which they are active ^142–144^. Eighth, the pairwise analysis mainly detects genes with high levels of expression, and thus genes with low expression in neurons would not be detected by our framework ^145,146^. Finally, our analysis relies on sequencing data for validation, which remains underpowered to implicate most genes.

### Reduced ATP2B2 expression reflects broader postsynaptic density dysregulation

Given *ATP2B2’s* role in synaptic calcium dynamics, we hypothesized that its reduced expression reflects broader disruption of glutamatergic neuron biology in schizophrenia. We tested this using a latent-factor analysis of synaptic gene programs (SNAP) ^23^., regressing gene expression on SNAP-derived donor loadings in dorsolateral prefrontal cortex neurons ( See Methods). We identified 1,277 genes with significant and positive associations in glutamatergic neurons (FDR < 5%) that were not similarly associated in GABAergic neurons (FDR < 5%). Gene set enrichment analysis^147^ of these genes revealed strong overrepresentation of postsynaptic density organization. Notably, several top-ranked genes implicated in autism and neurodevelopmental disorders, including *KCNB1, KCNMA1, SETBP1*, and *GABRB3*, were also specifically downregulated in glutamatergic neurons from schizophrenia donors. Together, these findings suggest a broader perturbation of postsynaptic density components, of which ATP2B2 is a functional constituent, pointing to coordinated dysregulation of calcium-handling and signaling machinery at excitatory synapses.

## Supplementary methods

### SNAP analysis

SNAP (Synaptic Neuron and Astrocyte Program) is a concerted gene-expression program in which cortical neurons and astrocytes co-regulate synaptic genes, and which is reduced in both schizophrenia and ageing. To identify all genes with correlated expression behavior to *ATP2B2* within the glutamatergic neuronal component of this program, we utilized the matrix of gene-and donor-level loadings onto the SNAP-neuron component (SNAP-n) provided in ^148^. We performed a genome-wide regression analysis in which the scaled expression of each gene was regressed against the SNAP-n donor loading, adjusting for relevant biological and technical covariates including age, postmortem interval (PMI), and sex

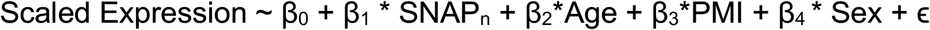

To mitigate bias from genes with high absolute expression but limited inter-individual variability, we used scaled expression values, ensuring that genes with extreme expression magnitudes do not disproportionately influence regression coefficients. This normalization was applied independently to both glutamatergic and GABAergic neuronal populations. We then restricted our analysis to genes showing statistically significant associations with SNAP-n donor loading in glutamatergic neurons while remaining non-significant in GABAergic neurons, thereby isolating genes with correlated expression behavior to *ATP2B2* in a cell-type-specific manner (Figure 12d)

### Gene Enrichment

For each trait with a GWAS and a corresponding exome-sequencing data, we define enrichment as the mean prioritization score (TauSum) for the nominated genes passing a significance threshold, divided by the scores of genes below that threshold. Given the heterogeneity in power between different neuropsychiatric traits (e.g;developmental disorders versus bipolar disorders), we maximized power by taking different thresholds. In the case of well powered sequencing data, we used FDR significant genes for Autism Spectrum Disorder and neurodevelopmental disorders, as per the original publications (FDR < 0.001 for ASD^72^ leading to 72 genes, 1.3e-06 for NDD^38^). For schizophrenia, bipolar disorder and epilepsy, we used a threshold of p < 0.01.

### Bootstrapping

Resampling methods were used to evaluate the robustness of gene prioritization scores for genes implicated in neuropsychiatric phenotypes (Figure 2a). Specifically, 10,000 bootstrap resamples were generated to estimate the distribution of prioritization scores within two gene sets: those designated as significant and those deemed not significant in the corresponding exome sequencing study. For each iteration, genes from the significant and not-significant groups were sampled with replacement to compute the mean prioritization score. This procedure generated empirical distributions of mean scores for each group, from which 95% confidence intervals were derived. The lower and upper bounds of these intervals were then compared between groups, using the ratio of the lower bound of one group to the upper bound of the other as a measure of separation between their score distributions.

### Voltage-gated gene set analysis

For the analysis of ion channels (Figure 3), we used the HUGO Gene Nomenclature Committee (HGNC) gene group report for Ion channels^54^, containing a total of 330 genes within the group and classification of ion channels by either gating mechanism (e.g voltage or ligand based) or by major permeant ion (channel type). At the first stage, we hierarchically tested the counts of ultra-rare PTV and MPC > 3 variants in genes partitioned into the two major non-overlapping subdivisions within this group (first being channel type and gating mechanism, and then by ligand-gated and voltage-gated). We filter out genes that are not expressed in the brain, defined as genes that have median transcripts per million (TPM) < 1 from GTEx^149^. Given the enrichment of Ca^2+^ genes in our analysis as well as previous implications of these genes in exome and GWAS studies, we focused on voltage-gated ion channels in our central analysis.

### Enrichment of prioritized genes from previous GWAS

For major depression, we selected prioritized genes as high-confidence genes as described in the original study^150^. For PTSD, the Tiers are based on the criterion established in ^151^, whereby the likelihood of being the causal risk gene) and tier 2 (prioritized over other GWAS-implicated genes, but lower likelihood than tier 1 of being the causal gene). For IQ^9^, we used the list of genes from their prioritized analysis. For epilepsy, we used all the genes prioritized as described in the original study and shown in the manhattan plot^152^. For schizophrenia, we tested against the Finemap set as well as the SMR set, while we tested against the SMR set for bipolar disorder.

### Case-control SCHEMA analysis

For schizophrenia, we downloaded all missense variants from the SCHEMA browser (https://schema.broadinstitute.org/) and downloaded all variants for *ATP2B2* (ENSG ID: ENSG00000157087), included those not used in the original analysis and generated scores in AlphaMissense and MisFit *s*.

### Mutagenesis primers for ATP2B2 constructs

- D454N forward primer: 5’-TGCCACAGCTATTTGCTCT**A**ATAAGACAGGTACTTTGAC-3’
- D454N reverse primer: 5’-GTCAAAGTACCTGTCTTAT**T**AGAGCAAATAGCTGTGGCA-3’
- T107M forward primer: 5’-CATTGCAAGATGTTA**TG**TTGATCATATTAGAAATCG-3’
- T107M reverse primer: 5’-CTTCCCATACCAATTGTAAAAATGTCTTTGG-3’
- E412K forward primer: 5’-CGCTGTACCT**A**AAGGTTTACCAT-3’
- E412K reverse primer: 5’-ACAACTAATACAGTGACACC-3’
- P415S forward primer: 5’-CTGAAGGTTTA**T**CATTGGCAGTTACAATTTC-3’
- P415S reverse primer: 5’-GTACAGCGACAACTAATACAGTGACAC-3’
- M684V forward primer: 5’-GTATCACCGTAAGA**G**TGGTTACTGG-3’
- M684V reverse primer: 5’-CTGCTCTTTGACATTTTCTTATAGCTTCTG-3’
- D688N forward primer: 5’-GTAAGAATGGTTACTGGT**A**ACAATATTAACACTG-3’
- D688N reverse primer: 5’-GGTGATACCTGCTCTTTGACATTTTC-3’
- G776R forward primer: 5’-CAGTTACCGGTGAT**C**GTACTAATGAC-3’
- G776R reverse primer: 5’-CGACAACTTGTCTTTGTTCAGTGTG-3’

## Acknowledgements

We thank Jacob Ulirsch, Tushar Kamath, Kaitlin Samocha, Beryl Cummings and members of the McCarroll laboratory including Marta Florio, Steven Burger, Yong Hoon Kim, Avin Veerakumar, as well as members of the Stanley Center including Ed Scolnick, Alkes Price, Alexander Gusev and Benjamin Neale for helpful discussions. and the brain tissue donors and their families, without whom this study would not be possible.

## Contributions

S.G, S.A.M and M.J.D designed the study. S.G and M.J.D performed the single-cell analysis with input from E.L., M.G., T.S., N.K. and S.A.M. J.Y and N.K. performed the synaptic proteomic analysis of ATP2B2. N.C.S, P.N and C.S designed and tested ATP2B2 variants in the colorimetric assay. M.A.B designed the cellular assay to test ATP2B2 variants with input from S.G, M.J.D, J.W. and J.Q.P. M.A.B and R.L performed the assay. F.K.S collected ATP2B2 variants for the ASD/NDD analysis. H.F. and S.G performed the 3D neighborhood enrichment analysis with input from M.J.D.

## Data availability

The Open Targets^115^ data were downloaded from https://www.opentargets.org/.

### The Autism Sequencing Consortium (ASC)

Branko Aleksic^42^, Mykyta Artomov^7,8,11,43^, Mafalda Barbosa^3,6^, Elisa Benetti^44,45^, Catalina Betancur^41^, Monica Biscaldi-Schafer^46^, Anders D. Børglum^47,48,49,50^, Harrison Brand^7,11,43,51^, Alfredo Brusco^52,53^, Joseph D. Buxbaum^1,2,3,4,5,6^, Gabriele S. Campos^14^, Simona Cardaropoli^54^, Diana Carli^54^, Angel Carracedo^55,56^, Marcus C. Y. Chan^57^, Andreas G. Chiocchetti^42^, Brian H. Y. Chung^57^, Brett Collins^1,2,6^, Ryan L. Collins^7,11,33,43^, Edwin H. Cook^38^, Hilary Coon^58,59^, Claudia I. S. Costa^14^, Michael L. Cuccaro^17,18^, David J. Cutler^60^, Mark J. Daly^7,8,9,10,34,35^, Silvia De Rubeis^1,2,4,6,29^, Bernie Devlin^12^, Ryan N. Doan^61^, Enrico Domenici^62^, Shan Dong^63^, Chiara Fallerini^44,45^, Magdalena Fernandez^16^, Montserrat Fernández-Prieto^55,64^, Giovanni Battista Ferrero^54^, Eugenio Ferro^16^, Jennifer Foss-Feig^1,2,6^, Christine M. Freitag^42^, Jack M. Fu^7,10,11^, Liliana Galeano^19^, J. Jay Gargus^65^, Sherif Gerges^7,8,11,43^, Elisa Giorgio^52^, Ana Cristina D. E. S. Girardi^14^, Stephen Guter^66^, Emily Hansen-Kiss^67^, Erina Hara^1,2^, Gail E. Herman^68^, Luis C. Hernandez^19^, Irva Hertz-Picciotto^20^, David M. Hougaard^46,69^, Christina M. Hultman^70^, Suma Jacob^66^, Miia Kaartinen^71^, Lambertus Klei^12^, Alexander Kolevzon^1,2,30^, Itaru Kushima^47,72^, Maria C. Lattig^19^, So Lun Lee^57^, Terho Lehtimäki^73^, Lindsay Liang^63^, Carla Lintas^74^, Alicia Ljungdahl^63^, Andrea del Pilar Lopez^15^, Caterina Lo Rizzo^44,45^, Yunin Ludena^20^, Patricia Maciel^75^, Behrang Mahjani^1,2,3,6,36,37^, Nell Maltman^66^, Marianna Manara^45,76^, Dara S. Manoach^77^, Gal Meiri^78,79^, Idan Menashe^80,81^, Judith Miller^82,83^, Nancy Minshew^12^, Matthew Mosconi^84^, Marina Natividad Avila^1,2,3,4,5,6^, Rachel Nguyen^65^, Norio Ozaki^47,85^, Aarno Palotie^7,9,35,86^, Mara Parellada^87^, Maria Rita Passos-Bueno^14^, Lisa Pavinato^52^, Katherine P. Peña^19^, Minshi Peng^88^, Margaret Pericak-Vance^17,18^, Antonio M. Persico^89^, Isaac N. Pessah^20^, Thariana Pichardo^1,2,4,6^, Kaija Puura^71^, Abraham Reichenberg^1,2,6,90^, Alessandra Renieri^44,45,76^, Kathryn Roeder^39,40^, Catherine Sancimino^1,2^, Stephan J. Sanders^91,92,93^, Sven Sandin^1,2,70^, F. Kyle Satterstrom^7,8,9^, Stephen W. Scherer^94,95^, Sabine Schlitt^42^, Rebecca J. Schmidt^20^, Lauren Schmitt^66^, Katja Schneider-Momm^42^, Paige M. Siper^1,2,6^, Laura Sloofman^1,2,3,4,5,6^, Moyra Smith^65^, Renee Soufer^1,2^, Christine R. Stevens^7,8,9^, Pål Suren^96^, James S. Sutcliffe^97,98^, John A. Sweeney^99^, Michael E. Talkowski^7,8,10,11,33^, Flora Tassone^20,31^, Karoline Teufel^42^, Elisabetta Trabetti^100^, Slavica Trajkova^52^, M. Pilar Trelles^32^, Brie Wamsley^101^, Jaqueline Y. T. Wang^14^, Lauren A. Weiss^63^, Mullin H. C. Yu^57^, Ryan Yuen^94^

^1^Seaver Autism Center for Research and Treatment, Icahn School of Medicine at Mount Sinai, New York, New York, USA. ^2^Department of Psychiatry, Icahn School of Medicine at Mount Sinai, New York, New York, USA. ^3^Department of Genetics and Genomic Sciences, Icahn School of Medicine at Mount Sinai, New York, New York, USA. ^4^Friedman Brain Institute, Icahn School of Medicine at Mount Sinai, New York, New York, USA. ^5^Department of Neuroscience, Icahn School of Medicine at Mount Sinai, New York, New York, USA. ^6^The Mindich Child Health and Development Institute, Icahn School of Medicine at Mount Sinai, New York, New York, USA. ^7^Program in Medical and Population Genetics, Broad Institute of MIT and Harvard, Cambridge, Massachusetts, USA. ^8^Stanley Center for Psychiatric Research, Broad Institute of MIT and Harvard, Cambridge, Massachusetts, USA. ^9^Analytic and Translational Genetics Unit, Department of Medicine, Massachusetts General Hospital, Boston, Massachusetts, USA. ^10^Center for Genomic Medicine, Department of Medicine, Massachusetts General Hospital, Boston, Massachusetts, USA. ^11^Department of Neurology, Massachusetts General Hospital and Harvard Medical School, Boston, Massachusetts, USA. ^12^Department of Psychiatry, University of Pittsburgh School of Medicine, Pittsburgh, Pennsylvania, USA. ^13^Division of Research, Kaiser Permanente Northern, Pleasanton, California, USA. ^14^Centro de Estudos do Genoma Humano e Células-Tronco, Departamento de Genética e Biologia Evolutiva, Instituto de Biociências, Universidade de São Paulo, São Paulo, Brasil. ^15^Facultad de Medicina, Universidad de los Andes, Bogotá, Colombia. ^16^Instituto Colombiano del Sistema Nervioso, Clínica Montserrat, Bogotá, Colombia. ^17^John P. Hussman Institute for Human Genomics, University of Miami Miller School of Medicine, Miami, Florida, USA. ^18^The Dr. John T. Macdonald Foundation Department of Human Genetics, University of Miami Miller School of Medicine, Miami, Florida, USA. ^19^Facultad de Ciencias, Universidad de los Andes, Bogotá, Colombia. ^20^MIND (Medical Investigation of Neurodevelopmental Disorders) Institute, University of California Davis, Davis, California, USA. ^21^Department of Psychiatry, Yale University School of Medicine, New Haven, Connecticut, USA. ^22^National Center of Posttraumatic Stress Disorders, VA CT Healthcare Center, West Haven, Connecticut, USA. ^23^Centro Ann Sullivan del Peru, Lima, Peru. ^24^Center Ann Sullivan International, Lawrence, Kansas, USA. ^25^Hospital Psiquiátrico Infantil Dr. Juan N. Navarro, Ciudad de México, Mexico. ^26^Universidad Nacional Autónoma de México, Ciudad de México, Mexico. ^27^Kaiser Permanente School of Medicine, Pasadena, California, USA. ^28^Departamento de Genética, Subdirección de Investigaciones Clínicas, Instituto Nacional de Psiquiatría Ramón de la Fuente Muñiz México, Ciudad de México, Mexico. ^29^The Alper Center for Neural Development and Regeneration, Icahn School of Medicine at Mount Sinai, New York, New York, USA. ^30^Department of Pediatrics, Icahn School of Medicine at Mount Sinai, New York, New York, USA. ^31^Department of Biochemistry and Molecular Medicine, University of California Davis, School of Medicine, Davis, California, USA. ^32^Psychiatry and Behavioral Sciences, Boston Children’s Hospital, Boston, Massachusetts, USA. ^33^Program in Bioinformatics and Integrative Genomics, Harvard Medical School, Boston, Massachusetts, USA. ^34^Department of Medicine, Harvard Medical School, Boston, Massachusetts, USA. ^35^Institute for Molecular Medicine Finland (FIMM), University of Helsinki, Helsinki, Finland. ^36^Department of Artificial Intelligence and Human Health, Icahn School of Medicine at Mount Sinai, New York, New York, USA. ^37^Department of Molecular Medicine and Surgery, Karolinska Institutet, Stockholm. ^38^Department of Psychiatry, University of Illinois Chicago, Chicago, Illinois, USA. ^39^Department of Statistics, Carnegie Mellon University, Pittsburgh, Pennsylvania, USA. ^40^Computational Biology Department, Carnegie Mellon University, Pittsburgh, Pennsylvania, USA. ^41^Sorbonne Université, INSERM, CNRS, Institut de Biologie Paris Seine, Center for Neuroscience at Sorbonne Université, Paris, France. ^42^Department of Psychiatry, Graduate School of Medicine, Nagoya University, Nagoya, Japan. ^43^Center for Genomic Medicine, Massachusetts General Hospital, Boston, Massachusetts, USA. ^44^Med Biotech Hub and Competence Center, Department of Medical Biotechnologies, University of Siena, Siena, Italy. ^45^Medical Genetics, University of Siena, Siena, Italy. ^46^Department of Child and Adolescent Psychiatry, Psychosomatics and Psychotherapy, Goethe University Frankfurt, Frankfurt, Germany. ^47^The Lundbeck Foundation Initiative for Integrative Psychiatric Research, iPSYCH, Aarhus, Denmark. ^48^Department of Biomedicine—Human Genetics, Aarhus University, Aarhus, Denmark. ^49^Center for Genomics and Personalized Medicine, Aarhus, Denmark. ^50^Bioinformatics Research Centre, Aarhus University, Aarhus, Denmark. ^51^Pediatric Surgical Research Laboratories, Department of Surgery, Massachusetts General Hospital, Boston, Massachusetts, USA. ^52^Department of Medical Sciences, University of Torino, Turin, Italy. ^53^Medical Genetics Unit, ‘Città della Salute e della Scienza’ University Hospital, Turin, Italy. ^54^Department of Public Health and Pediatrics, University of Torino, Turin, Italy. ^55^Grupo de Medicina Xenómica, Centro de Investigación en Red de Enfermedades Raras (CIBERER), CIMUS, Universidade de Santiago de Compostela, Santiago de Compostela, Spain. ^56^Fundación Pública Galega de Medicina Xenómica, Servicio Galego de Saúde (SERGAS), Santiago de Compostela, Spain. ^57^Department of Pediatrics and Adolescent Medicine, Duchess of Kent Children’s Hospital, The University of Hong Kong, Hong Kong Special Administrative Region, China. ^58^Department of Internal Medicine, University of Utah, Salt Lake City, Utah, USA. ^59^Department of Psychiatry, Huntsman Mental Health Institute, University of Utah, Salt Lake City, Utah, USA. ^60^Department of Human Genetics, Emory University School of Medicine, Atlanta, Georgia, USA. ^61^Division of Genetics and Genomics, Boston Children’s Hospital, Boston, Massachusetts, USA. ^62^Department of Cellular, Computational and Integrative Biology, University of Trento, Trento, Italy. ^63^Department of Psychiatry, UCSF Weill Institute for Neurosciences, University of California San Francisco, San Francisco, California, USA. ^64^Neurogenetics group, Instituto de Investigación Sanitaria de Santiago (IDIS-SERGAS), Santiago de Compostela, Spain. ^65^Center for Autism Research and Translation, University of California Irvine, Irvine, California, USA. ^66^Institute for Juvenile Research, Department of Psychiatry, University of Illinois at Chicago, Chicago, Illinois, USA. ^67^Department of Diagnostic and Biomedical Sciences, University of Texas Health Science Center at Houston, School of Dentistry, Houston, Texas, USA. ^68^The Research Institute at Nationwide Children’s Hospital, Columbus, Ohio, USA. ^69^Center for Neonatal Screening, Department for Congenital Disorders, Statens Serum Institut, Copenhagen, Denmark. ^70^Department of Medical Epidemiology and Biostatistics, Karolinska Institutet, Stockholm, Sweden. ^71^Department of Child Psychiatry, Tampere University and Tampere University Hospital, Tampere, Finland. ^72^Medical Genomics Center, Nagoya University Hospital, Nagoya, Japan. ^73^Department of Clinical Chemistry, Fimlab Laboratories and Finnish Cardiovascular Research Center-Tampere, Faculty of Medicine and Health Technology, Tampere University, Tampere, Finland. ^74^Service for Neurodevelopmental Disorders, University Campus Bio-medico of Rome, Rome, Italy. ^75^Life and Health Sciences Research Institute, School of Medicine, University of Minho, Braga, Portugal. ^76^Genetica Medica, Azienda Ospedaliera Universitaria Senese, Siena, Italy. ^77^Department of Psychiatry, Massachusetts General Hospital and Harvard Medical School, Boston, Massachusetts, USA. ^78^The Azrieli National Center for Autism and Neurodevelopment Research, Ben-Gurion University of the Negev, Beer-Sheva, Israel. ^79^Pre-School Psychiatry Unit, Soroka University Medical Center, Beer Sheva, Israel. ^80^Department of Public Health, Ben-Gurion University of the Negev, Beer-Sheva, Israel. ^81^National Autism Research Center of Israel, Ben-Gurion University of the Negev, Beer-Sheva, Israel. ^82^Children’s Center for Autism Research and Training, University of Kansas, Lawrence, Kansas, USA. ^83^Department of Psychiatry, University of Utah, Salt Lake City, Utah, USA. ^84^Life Span Institute and Kansas Center for Autism Research and Training, University of Kansas, Lawrence, Kansas, USA. ^85^Institute for Glyco-core Research (iGCORE), Nagoya University, Nagoya, Japan. ^86^Psychiatric & Neurodevelopmental Genetics Unit, Department of Psychiatry, Massachusetts General Hospital, Boston, Massachusetts, USA. ^87^Department of Child and Adolescent Psychiatry, Hospital General Universitario Gregorio Marañón, IiSGM, CIBERSAM, School of Medicine Complutense University, Madrid, Spain. ^88^Department of Statistics and Data Science, Carnegie Mellon University, Pittsburgh, Pennsylvania, USA. ^89^Interdepartmental Program ‘Autism 0-90’, ‘Gaetano Martino’ University Hospital, University of Messina, Messina, Italy. ^90^Department of Environmental Medicine and Public Health, Icahn School of Medicine at Mount Sinai, New York, New York, USA. ^91^Institute of Developmental and Regenerative Medicine, Department of Paediatrics, University of Oxford, Oxford, UK ^92^Department of Psychiatry and Behavioral Sciences, UCSF Weill Institute for Neurosciences, University of California, San Francisco, San Francisco, California, USA. ^93^New York Genome Center, New York, New York, USA. ^94^Program in Genetics and Genome Biology, The Centre for Applied Genomics, The Hospital for Sick Children, Toronto, Ontario, Canada. ^95^Department of Molecular Genetics and McLaughlin Centre, University of Toronto, Toronto, Ontario, Canada. ^96^Norwegian Institute of Public Health, Oslo, Norway. ^97^Department of Molecular Physiology & Biophysics and Psychiatry, Vanderbilt University School of Medicine, Nashville, Tennessee, USA. ^98^Vanderbilt Genetics Institute, Vanderbilt University School of Medicine, Nashville, Tennessee, USA. ^99^Department of Psychiatry, University of Cincinnati, Cincinnati, Ohio, USA. ^100^Department of Neurosciences, Biomedicine and Movement Sciences, Section of Biology and Genetics, University of Verona, Verona, Italy. ^101^Program in Neurogenetics, Department of Neurology, David Geffen School of Medicine, University of California Los Angeles, Los Angeles, California, USA.

